# Transcriptional Responses of Cancer Cells to Heat Shock-Inducing Stimuli Involve Amplification of Robust HSF1 Binding

**DOI:** 10.1101/2022.12.14.519647

**Authors:** Sayantani Ghosh Dastidar, Bony De Kumar, Bo Lauckner, Damien Parrello, Danielle Perley, Maria Vlasenok, Antariksh Tyagi, Nii Koney-Kwaku Koney, Ata Abbas, Sergei Nechaev

## Abstract

Responses of cells to signals are increasingly discovered to involve the binding of sequence-specific transcription factors outside of known target genes. We wanted to determine to what extent the genome-wide binding and function of a transcription factor are shaped by the cell type versus the stimulus. To do so, we induced the Heat Shock Response pathway in two distant cell lines with two different stimuli and related the binding of its master regulator HSF1 to nascent RNA and chromatin accessibility. We show that HSF1 binding patterns robustly retain their identity under different magnitudes of activation so that common HSF1 binding is globally associated with stimulus-specific transcription outcomes. HSF1-induced increase in DNA accessibility was modest in scale but occurred predominantly at remote genomic sites. Apart from regulating transcription at existing elements including promoters and enhancers, responses to heat shock may directly engage inactive chromatin.

## INTRODUCTION

Transcriptional responses to stimuli involve the binding of sequence-specific factors to DNA to regulate their target genes. The Heat Shock Response (HSR) is an evolutionarily conserved cellular stress defense mechanism that can be triggered by heat and some other stimuli (1–4), and is frequently active in cancers (3, 5). Even though HSR is recognized to be a complex pathway involving numerous regulatory components (6–9), the Heat Shock Factor (HSF, HSF1 in mammals) is considered its master regulator. HSF1 is normally found in the cytoplasm (2, 10), but during stress is phosphorylated to translocate into the nucleus and bind to its cognate Heat Shock Response DNA Elements (HREs) (3, 11, 12). Foundational work in *Drosophila* demonstrated that HSR involves rapid appropriation of the RNA polymerase II (Pol II) machinery by a handful of Heat Shock Protein (HSP) genes, leading to their massive activation at the expense of the rest of the genome (13, 14). Binding to promoters of HSP genes to activate their transcription was presumed to be the function of HSF (15–17). This paradigm has shaped our current understanding of transcription regulation, particularly initiation and early elongation (10, 18, 19). However, recent studies revealed that the binding of HSF1 during HSR is not limited to HSP gene promoters, or to promoters at all, but also occurs at thousands of intergenic loci (6, 20). Moreover, HSF1 binding near promoters does not necessarily lead to transcription activation of nearby genes (6, 21). These findings challenge the established relationship between the binding of a transcription factor to its target sites and transcription.

Analysis of the human genome identifies over two hundred and eighty thousand HSF1 binding motifs in the hg19 genome assembly (22). However, profiling of HSF1 binding using Chromatin-Immunoprecipitation Sequencing (ChIP-seq) in human (3, 23) or mouse cells (6) finds on the order of ten thousand HSF1 peaks in a given dataset. This difference leaves ample room for HSF1 binding patterns to be flexible. We therefore asked to what extent the genome-wide binding of HSF1 during HSR varies by the cell type versus the stimulus and how this binding relates to nascent transcription. To answer these questions, we subjected two distant human cell lines, MCF-7 breast adenocarcinoma and K562 chronic myelogenous leukemia, to two different HSR-inducing stimuli: elevated temperature and arsenic, a toxic metalloid found in soil, air and water, at the ambient temperature (24). By following the binding of HSF1 along with Pol II, nascent transcription, the active promoter mark histone H3 lysine 4 trimethylation (H3K4Me3) and chromatin accessibility using the Assay for Transposase Accessible Chromatin with high-throughput sequencing (ATAC-seq) (25), we find that among several readouts including transcription, HSF1 signal exhibits the most salient genome-wide changes, which do not require the elevated temperature. HSR involves amplification of low-level HSF1 binding with genome-wide patterns that are distinct between cell lines but more consistent between stimuli. HSF1 binding provides a robust platform for context-specific transcription activation of promoters and enhancers as well as stimulus-dependent engagement of inactive chromatin sites.

## RESULTS

### Widespread HSF1 binding is a temperature-independent hallmark of heat shock response

We began by examining MCF7 breast adenocarcinoma cells during temperature-induced Heat Shock Response (HSR) (Fig. 1; Supplementary File 1; Supplementary Fig. 1). A 60-minute incubation at 42°C induced a control HSP gene seen as spreading of the Pol II and nascent RNA signal into the gene body (Fig. 1a). HSF1 also showed characteristic binding at the *HSPH1* gene promoter near the transcription start site (TSS) (Fig. 1a), similar to that widely observed on HSP genes (3, 6, 23). Genome-wide, ChIP-sequencing using a previously validated anti-HSF1 antibody (20) showed widespread HS-dependent signal at thousands of sites at (+/-1kb) and outside of annotated gene promoters (Fig. 1b, c), with ~18,000 peaks (q<0.01) identified between two independent biological replicates (Supplementary Fig. 2a; Supplementary File 2). A different anti-HSF1 antibody (3) showed 75% overlap in peak locations in MCF7 cells (Supplementary Fig. 2b). Genome-wide changes for other readouts, however, including transcription itself, were less pronounced. An active promoter histone mark H3K4me3 showed no drastic changes in HS compared to untreated control (non-heat shock, NHS) cells (Fig. 1a, d). There was even a modest decrease in H3K4me3 signal at the immediate promoter-proximal regions of highly activated genes including HSP70 *(HSPA1B)* (Fig. 1a; Supplementary Fig. 3), similar to earlier findings in yeast and *Drosophila* (26–29). Nascent RNA analysis using Precision nuclear Run-On sequencing (PRO-seq) showed 2,288 genes upregulated in HS (p-adj < 0.05) (Supplementary File 3). Pol II ChIP-seq and PRO-seq signal was retained at transcription initiation sites outside of activated promoters as well, consistent with ongoing transcription in HS outside of activated genes genome-wide (Fig. 1a, e; Supplementary Fig. 3, 4) (6, 30).

**Fig. 1:**
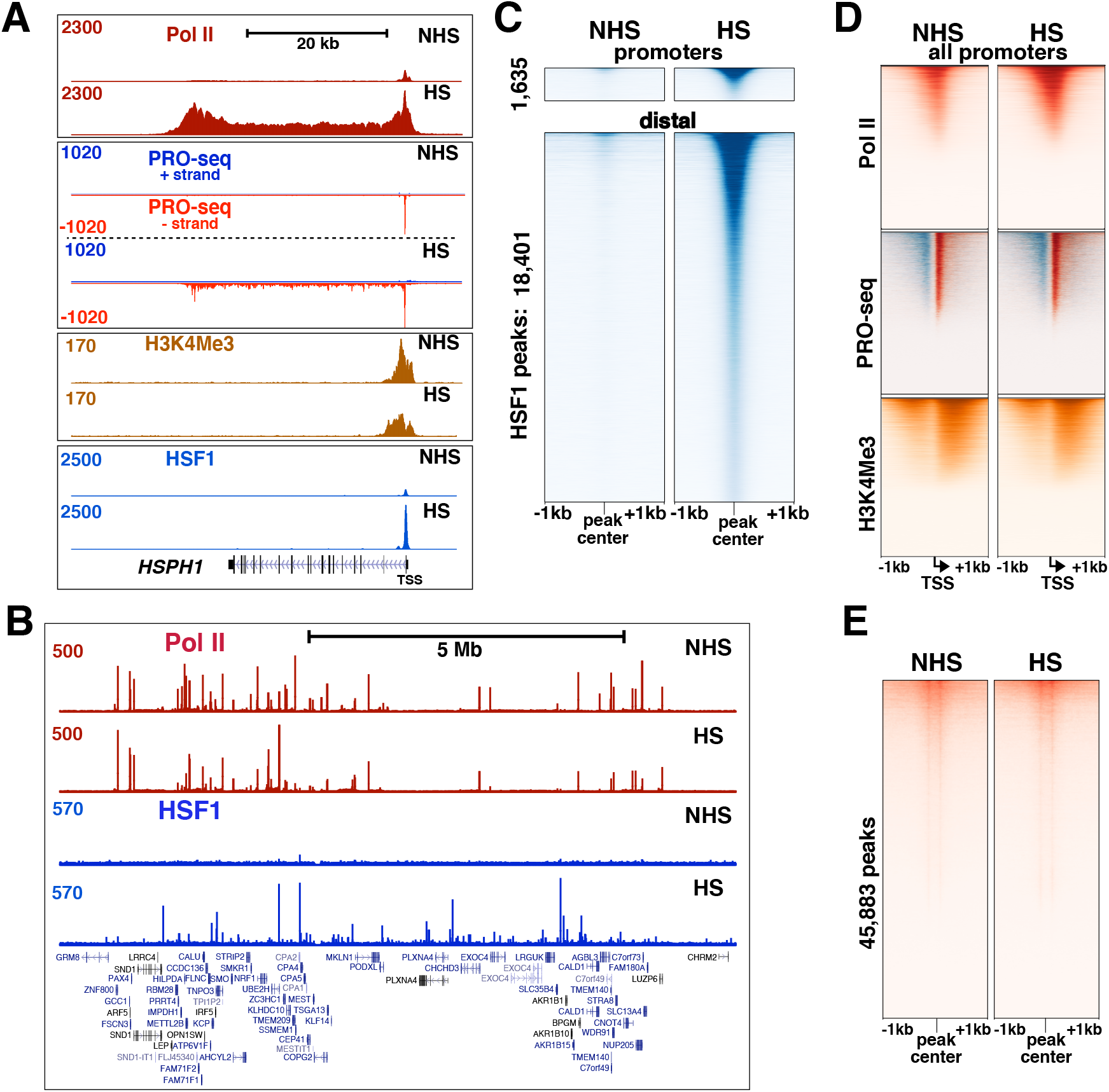
Genome-wide response to HS. **a.** UCSC genome browser tracks of ChIP-seq and PRO-seq datasets for MCF7 cells in control, non heat shock (NHS) and heat shock (HS) conditions are shown on a HS-inducible gene. PRO-seq tracks are separately for plus and minus strands. The gene TSS is indicated on the right. **b.** Pol II and HSF1 ChIP-seq tracks for NHS and HS conditions showing a randomly selected ~10Mb section of the genome, with genes indicated underneath. Some duplicate genes were removed from the image. **c.** Heatmaps showing HSF1 ChIP-seq signal centered at HSF1 peaks in promoter-proximal and distal regions in NHS and HS conditions, and sorted by HS peak intensity. **d.** Heatmaps showing the indicated readouts centered at TSSs for all 23K promoters. PRO-seq signal is shown in red for the sense strand signal and in blue for signal antisense with respect to genes. **e.** Heat-map of PRO-seq signal centered around PRO-seq peaks located outside of gene regions with respect to the plus strand of the genome. The data indicate prevalent pausing at positions on either side approximately 50nt from the peak center. Heatmaps in (c-e) are sorted by signal intensity in HS samples and all datasets are scaled to their sequencing depths.

Upregulation of *HSP* genes has been previously induced at the ambient temperature in the presence of inorganic arsenic (24). This prompted us to ask whether genome-wide HSF1 binding would also occur in the absence of heat. Treatment of MCF-7 cells with sodium meta-arsenite (As) at 37°C showed transcriptional activation of control HS genes, with the timing and magnitude of mRNA level increase resembling those induced by heat (Fig. 2a; Supplementary Fig. 5). Like HS, As induced characteristic HSF1 and Pol II binding at *HSPA1B* gene promoter as observed by ChIP-qPCR (Fig. 2b). ChIP-sequencing of As-treated MCF7 cells revealed widespread HSF1 binding (Fig. 2c, d; Supplementary File 2). HSF1 signal in each treatment at promoters was comparable in intensity although statistically higher than the signal at distant regions (p<0.0001) (Fig. 2e), with the intensity of HSF1 signal being comparable between HS and As treatments (Supplementary Fig. 6). Elevated temperature is therefore not a prerequisite for genome-wide HSF1 binding.

**Fig. 2:**
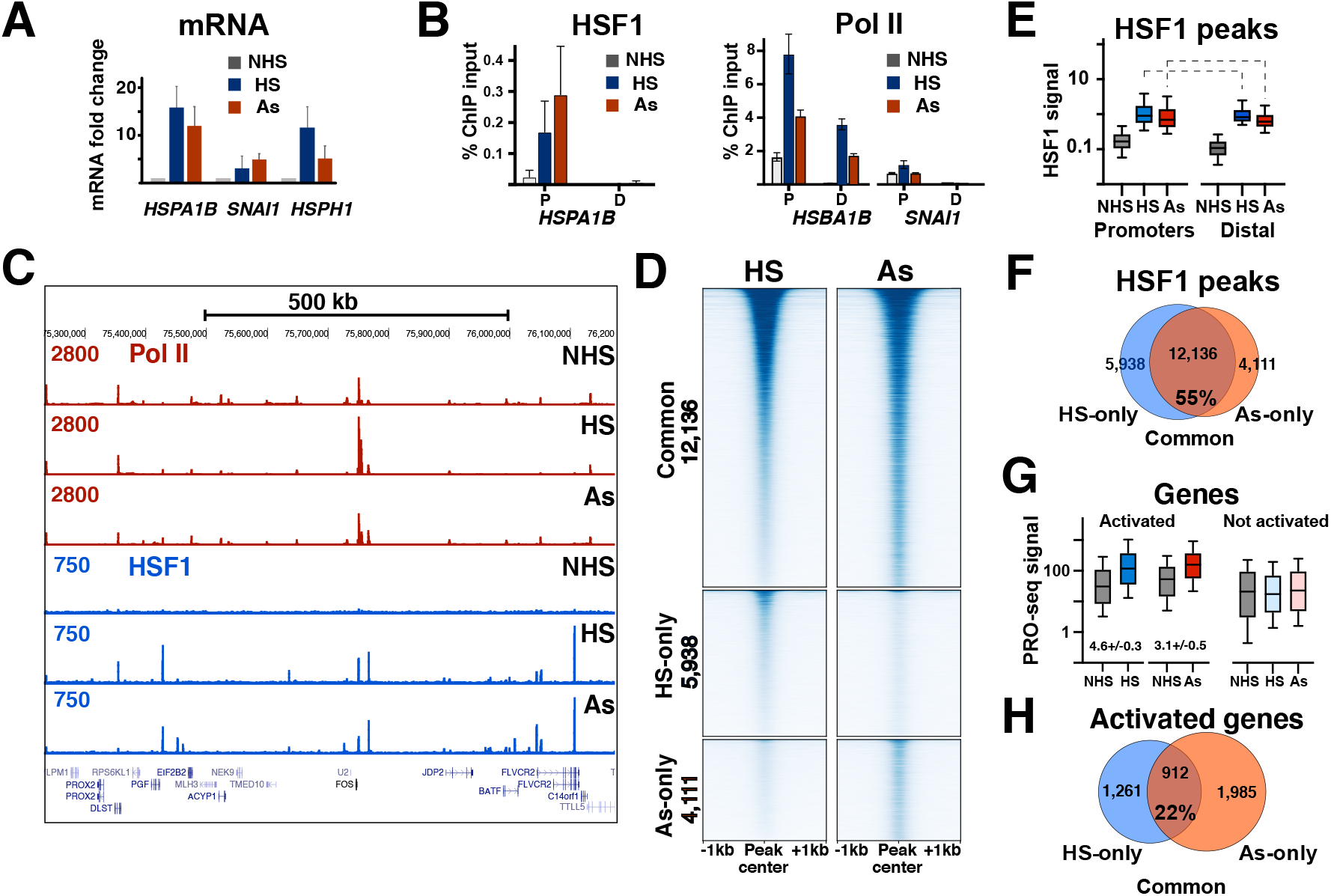
Widespread HSF1 binding is independent of elevated temperature. **a.** RT-qPCR showing fold change in HSP70, SNAI1, and HSPH1 mRNA transcripts compared to GAPDH in NHS, HS, and As conditions. **b**. ChIP-qPCR against HSF1 (left) and RNA Pol II (right) showing fold enrichment over input at promoter and distal regions of HSPA1B and SNAI1 genes in NHS, HS, and As conditions. **c.** UCSC genome browser tracks showing ChIP-seq signal from RNA Pol II and HSF1 on a ~1MB portion of the genome surrounding FOS gene that is activated in HS. **d.** Heat-map of peak-centered MCF7 HSF1 signal in HS and As conditions for peaks common for HS and As (top), exclusive for HS (middle) and exclusive for As (bottom). Sample assignment is color-coded on the right. **e.** Averaged HSF1 signal (Counts Per Million uniquely mapped reads) from peaks at promoters (TSS +/-1000) (1,695 peaks for HS and 2,079 for As) or distal regions (16,461 for HS and 14,305 for As). Promoter signal was higher for both As replicates (p<0.001) and for both HS replicates (p<0.045, p<0.0001) based on the Mann-Whitney test (comparisons shown with dashed lines). HS and As signal was not consistently higher between replicates (Supplementary Fig. 6). **f.** Venn diagram showing the numbers of common and exclusive MCF7 HSF1 peaks in HS and As conditions. The percentage of common peaks among all peaks is shown underneath. **g**. PRO-seq gene body signal density for activated genes (2,288 for HS and 2,974 for As) (left) and not activated genes (21,410 for HS and 20,723 for As) (right) in MCF7 cells (f). Data for this graph were normalized by sequencing coverage following rRNA removal. Numbers indicate the mean fold activation compared to the same genes in untreated cells, with the range between two biological replicates. **h**. Venn diagram showing common and condition-exclusive activated genes between HS and As treatments. The percentage of HS and As commonly activated among all activated is shown underneath.

Comparing HSF1 binding locations between HS and As treatments, a majority of peaks in MCF7 cells overlapped between them (Fig. 2f). Accordingly, sorting all HSF1 peaks by the signal intensity in one treatment retained their ordering in the other (Fig. 2d). All groups of peaks were dominated by the cognate HRE DNA sequence motifs (Supplementary Fig. 7). Examining nascent transcription in PRO-seq datasets, genes commonly activated in HS and As were enriched in stimulus response categories including classic *HSP* genes (Supplementary File 4). The fold gene activation was overall modestly higher in HS than As treatments in our hands (Fig. 2g), with As-induced transcription reaching saturation in titration experiments on control genes (Supplementary Fig. 5). Comparing HSF1 binding and gene activation, HSF1 peaks showed a higher overlap between treatments than did transcriptionally activated genes (Fig. 2f, h), as only about a fifth of activated genes were in common between treatments. This relationship persisted under increased stringency of calling HSF1 peaks or activated genes (Supplementary Fig. 8). Taken together, widespread HSF1 binding is a temperature-independent hallmark of HSR whose genome-wide patterns are more stable between stimuli than transcription.

### Common HSF1 binding and variable transcription activation

Two treatments inducing widespread HSF1 binding allowed us to probe the relationship between HSF1 promoter binding and gene transcription. Approximately 8% of gene promoters contained HSF1 peaks (Fig. 3a). For activated genes, the fraction of promoters with HSF1 peaks was at least 2-fold higher (Fig. 3b), broadly implicating HSF1 promoter binding in gene activation. However, most genes in both treatments in MCF7 cells were activated in the absence of HSF1 binding (Fig. 3b). Accordingly, most HSF1 binding at promoters was not associated with gene activation (Fig. 3c). These observations are consistent with previous work in HS (6, 21) and reinforce the notion of the overall disconnect between HSF1 binding and nearby gene transcription. To gain insight into possible reasons behind this disconnect, we compared HSF1 binding on activated versus not activated genes. While HSF1 peaks did show higher HSF1 signal at promoters of activated than not activated genes (Fig. 3d), transcription fold-activation was similar for genes activated with and without HSF1 (Fig. 3e). The difference between HSF1-bound and unbound promoters was, surprisingly, in basal activity, which was higher in NHS cells for genes whose promoters would bind HSF1 in HSR than those that do not. This held true for activated (Fig. 3e) and not activated genes alike (Fig. 3f). HSF1 thus favors basally active promoters regardless of whether the genes are activated in HSR or not.

**Fig. 3:**
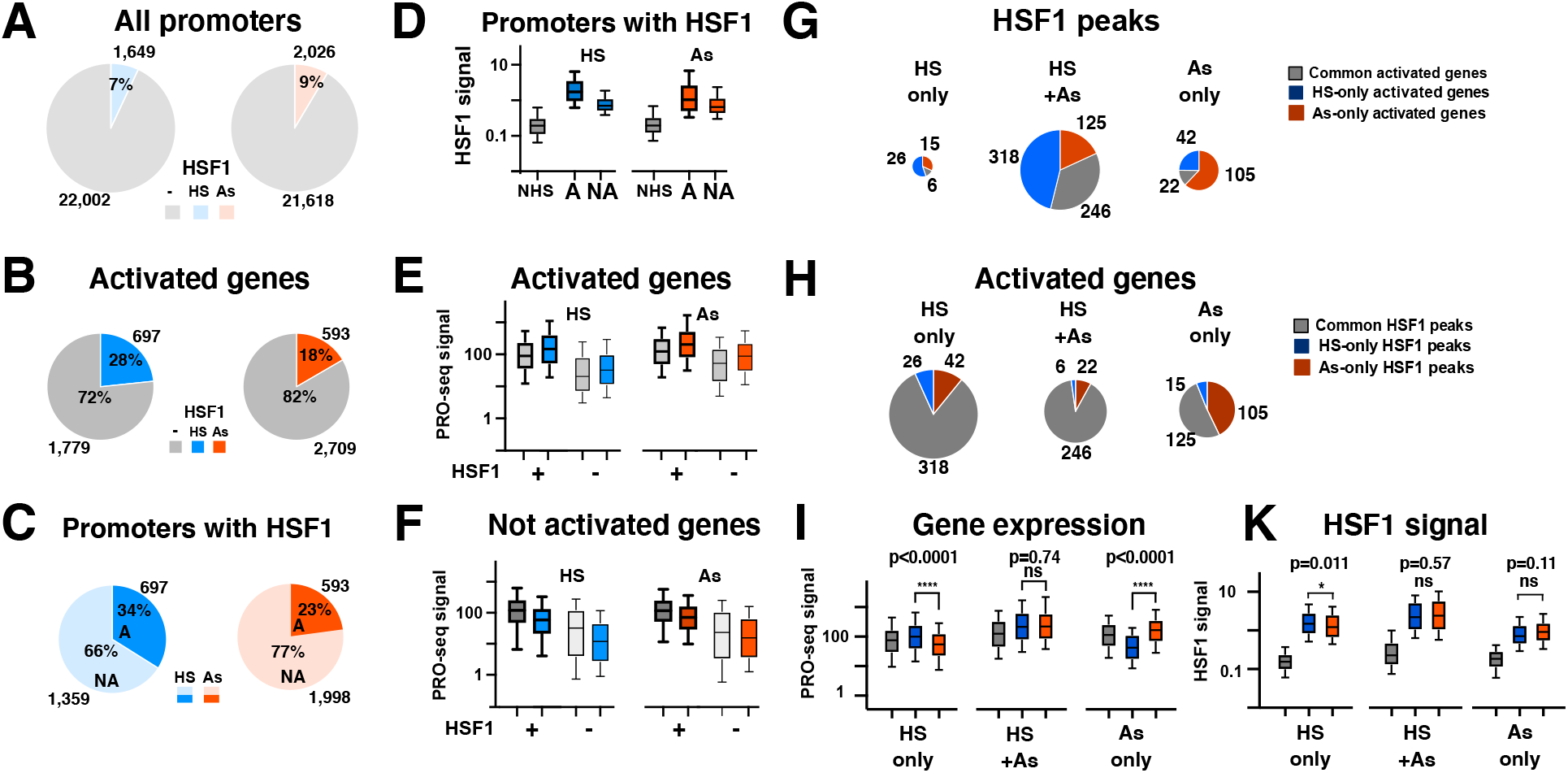
HSF1 binding at gene promoters is decoupled from transcriptional activation. Pie charts showing the fraction of promoters with or without HSF1 peaks within +/1kb interval for all 23K promoters (**a**) and promoters of activated genes (**b**). **c.** Pie chart showing the numbers and proportion of promoters containing HSF1 within +/-1kb interval for genes that are transcriptionally activated in the same treatment (darker color) or not activated (lighter color). HS is indicated in blue and As in red. **d.** Average HSF1 peak signal for genes containing HSF1 peaks at promoters as in (c) for activated (A) and not activated (NA) genes. NHS shows HSF1 signal for A and NA genes grouped together, in NHS conditions. Box plots show the mean with 10-90% confidence interval unless otherwise indicated. **e.** Gene expression based on PRO-seq gene body count density for activated genes with (+) (697 and 593 genes for HS and As, respectively) and without (-) HSF1 peaks at promoters, +/-1kb from the gene TSS (1,779 and 2,709 genes for HS and As). **f.** Gene expression for not activated genes with (+) (1,359 and 1,998 for HS and As, respectively) and without (-) HSF1 at promoters (20,223 for HS and 18,909 for As). **g.** Distribution of conditionally activated genes with HSF1 promoter peaks among those containing common and condition-unique HSF1 peaks. Each HSF1 peak category is indicated in a pie chart scaled to the number of genes in each, with slices indicating common (grey) or condition-specific activated genes (blue for HS-specific and red for As specific). **h.** Distribution of HSF1 peaks among genes common and uniquely activated in each treatment as in (g). Slices indicate common (grey) or condition-unique (blue for HS and red for As) HSF1 peaks at promoters. **i.** Gene expression based on PRO-seq read density (i) for gene groups in (h). **k.** HSF1 peak signal at NHS, HS, and As conditions for the genes shown in (h) and (i). Statistical significance is calculated based on p-value from Mann-Whitney test.

Next, we examined a subset of ~900 HSF1 peak regions that were associated with promoters of genes activated in either treatment. A vast majority of these HSF1 peaks were in common between treatments (Fig. 3g). HSF1 peaks unique to each treatment were relatively few, but nevertheless were enriched in genes activated in the same treatment, both for HS and As (Fig. 3g, left, right). Condition-specific HSF1 binding is therefore associated with gene activation in the same treatment. Pivoting the data to view activated genes, we noted no enrichment of HS-specific HSF1 peaks at genes activated in HS (Fig. 3h, left, middle). We did note that 43% of As-exclusive activated genes were associated with As-exclusive HSF1 promoter peaks (Fig. 3h, right), indicating that HSF1 can bind and potentially function at new sites outside of a generic response to temperature. However, As-specific activation involved only a modest number of genes. A vast majority of activated genes were associated with common HSF1 binding (Fig. 3h). Despite the differences in transcriptional outcomes, HSF1-bound genes activated only in HS- or only in As treatments did not show significant differences in HSF1 binding intensity by stringent criteria (p=0.011) (Fig. 3i, k) and no difference in binding positions (Supplementary Fig. 9). Taken together, comparison of two HSR-inducing stimuli points to widespread decoupling of HSF1 binding and gene activation wherein context-specific transcription is predominantly associated with common HSF1 binding.

### Distinct HSF1 patterns between cell lines

Having compared HSR in MCF7 cells induced with HS or As, we applied the same two stimuli to cells of an unrelated origin. K562 is a Tier I ENCODE leukemia cell line that has been previously examined for rapid responses to heat (23). RT-qPCR and PRO-seq showed that both MCF7 and K562 cells responded to HS or As by upregulating control genes (Fig. 4a, b; Fig. 2a). Genome-wide, HS treatment of K562 and MCF7 cells activated over a thousand genes (1,760 and 2,288, respectively) (Supplementary File 3), with 70% of upregulated genes from an existing K562 HS dataset (546 out of 778 genes) being in common with our K562 HS treatment (30). Either HS or As induced widespread HSF1 binding in both cell lines (Fig. 4b, e, Supplementary File 2), indicating that temperature independence of genome-wide HSF1 binding is not confined to a particular cell type.

**Fig. 4:**
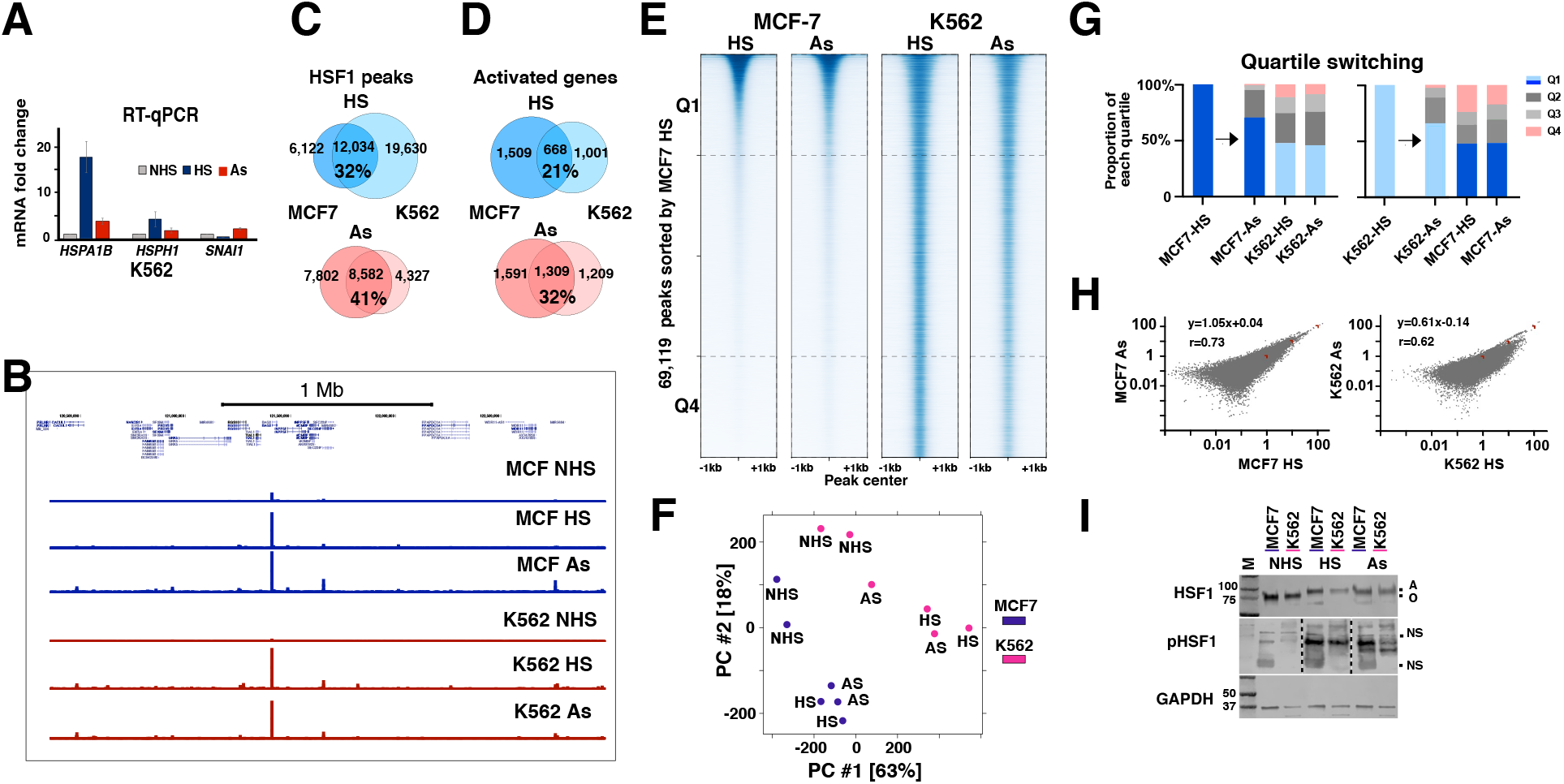
HSF1 follows cell type specific patterns of genome-wide binding. **a.** RT-qPCR showing fold-change in the indicated mRNA compared to GAPDH in K562 cells. **b.** UCSC browser tracks showing MCF7 (blue) and K562 (red) HSF1 ChIP-seq from a randomly selected ~3Mb portion of the genome. **c.** Venn diagram showing HSF1 peaks in HS (blue) and As (red) conditions between MCF7 and K562 cells. The percentage of common peaks among all peaks is shown underneath. **d.** Venn diagram showing activated genes in HS (blue) and As (red) conditions in MCF7 and K562 cells. The percentage of commonly activated genes among all activated genes is shown underneath. **e.** Heat-map showing HSF1 peak signal (+/- 1000bp from peak center) in MCF7 and K562 cells. Heatmap is sorted by MCF7 HS signal and is divided into 4 quartiles (Q1-Q4); each quartile contains approximately 17, 280 peaks), with top and bottom quartile boundaries indicated by dashed lines. **f.** PCA plot of HSF1 peaks in MCF7 (purple) versus K562 (pink) cells shown for each condition per replicate. **g.** Bar graph showing the changes of HSF1 peaks between quartiles. Distribution of MCF7 HS quartile 1 (Q1) peaks (left, dark blue) and K562 HS Q1 peaks (right, light blue) is shown across each quartile for all samples in (e) as indicated below the graph. **h.** Dot plots for HSF1 peak signal intensity between indicated treatments. Linear regression was calculated based on these datasets normalized by the sequencing depth, with Spearman correlation coefficient indicated. **i.** Western Blot showing Hsf1 and phosphorylated HSF1 (pHSF1) in NHS, HS and As conditions in MCF7 and K562 cells. GAPDH is used as a loading control. NS represents non-specific bands. A and O represent active (phosphorylated) and inactive forms of Hsf1 distinguished by different mobility on SDS gels. Numbers indicate molecular weight of marker bands (not visible in pHSF1 blots).

The overlap in HSF1 peak locations between the two cell lines was lower than that between HS and As treatments within a cell line (Fig. 4b; Fig. 2f). This is consistent with cell type specificity of HSF1 binding. However, the overlaps between HSF1 peaks were still higher than between activated genes (Fig. 4c, d), consistent with higher conservation of HSF1 binding and flexibility of transcription responses. Comparing HSF1 signal across all datapoints used in this work showed clear rank-ordering by the cell type rather than treatment (Fig. 4e) (3). By Principal Component Analysis (PCA), individual datasets separated by the cell type as well (Fig. 4f) (31). This separation was preserved even after HSF1 signal was normalized to DNA genomic copy number data for each cell line (32) or when only a subset of high confidence common peaks was used (Supplementary Fig. 10). Differences in HSF1 binding intensity among detected peaks can thus account for cell type specific binding patterns. To find sites with the most dramatic changes, we selected HSF1 peaks found in the opposite quartiles (Q1 and Q4) between datapoints when sorted by signal intensity. About 25% of the top quartile (Q1) MCF HS HSF1 peaks were found in the bottom quartile (Q4) of K562 HS peaks, compared to under 4% in Q4 of MCF7 As peaks (Fig. 4e, g). While we could not fully exclude the contribution of genetic mutations to differences in HSF1 binding, variations in genomic sequences based on available genomic information (32, 33) did not affect the prevalence of the HSF1 consensus sequence under these peaks (Supplementary Fig. 11). These changed peaks were predominantly intergenic, with their fraction at promoters (4.9%) being even lower than the overall HSF1 promoter peak fraction of 7% (Fig. 3a). Altogether, data indicate that HSF1 shows cell-type specificity in binding locations and intensity.

The observed cell type specificity of HSF1 binding may not be surprising as similar observations have been made in earlier studies (31). However, natural differences in stimulus intensity or cellular environment may induce responses at different magnitudes and therefore scramble the transcription factor binding patterns. K562 cells consistently exhibited a lower magnitude of HSF1 binding in response to As than to HS in our hands (Fig. 4h). While we did not investigate the underlying reasons, this observation is mirrored by lower levels of the active phosphorylated form of HSF1 in K562 cells treated with As compared to other datapoints (Fig. 4i). That K562 cells treated with As clustered with HS despite the lower overall signal indicates that genome-wide HSF1 patterns retain their identity under different magnitudes of HSR activation.

### Modest connection between HSF1 binding and nascent transcription extends beyond promoters

Even though over 80% of all HSF1 peaks were found in the intergenic regions, their density was several-fold higher at gene promoters despite the lack of enrichment of HRE sequence elements (Fig. 5a). Given that most HSF1 binding at promoters did not activate transcription (Fig. 3c) (6), we asked to what extent the sites where HSF1 binding is associated with gene activation are conserved between distant cell lines. HSF1 showed higher enrichment at promoters of genes activated in both cell lines than at those activated in only one, with about half of promoters for genes commonly activated between the cell lines containing HSF1 peaks (Fig. 5b). These commonly activated genes included the known heat shock response genes harboring peaks with the highest signal (Supplementary File 5). However, this group constitutes only a small fraction of genes activated between these two cell lines (Fig. 5b; Fig. 4d). Despite the higher signal (Supplementary Fig. 12), common HSF1 peaks barely showed a higher proportion at promoters when viewed genome-wide, effectively diluting the contribution from these conserved genes (Fig. 5c). This observation is consistent with a limited direct role of HSF1 in gene activation genome-wide apart from classic HSP genes.

**Fig. 5:**
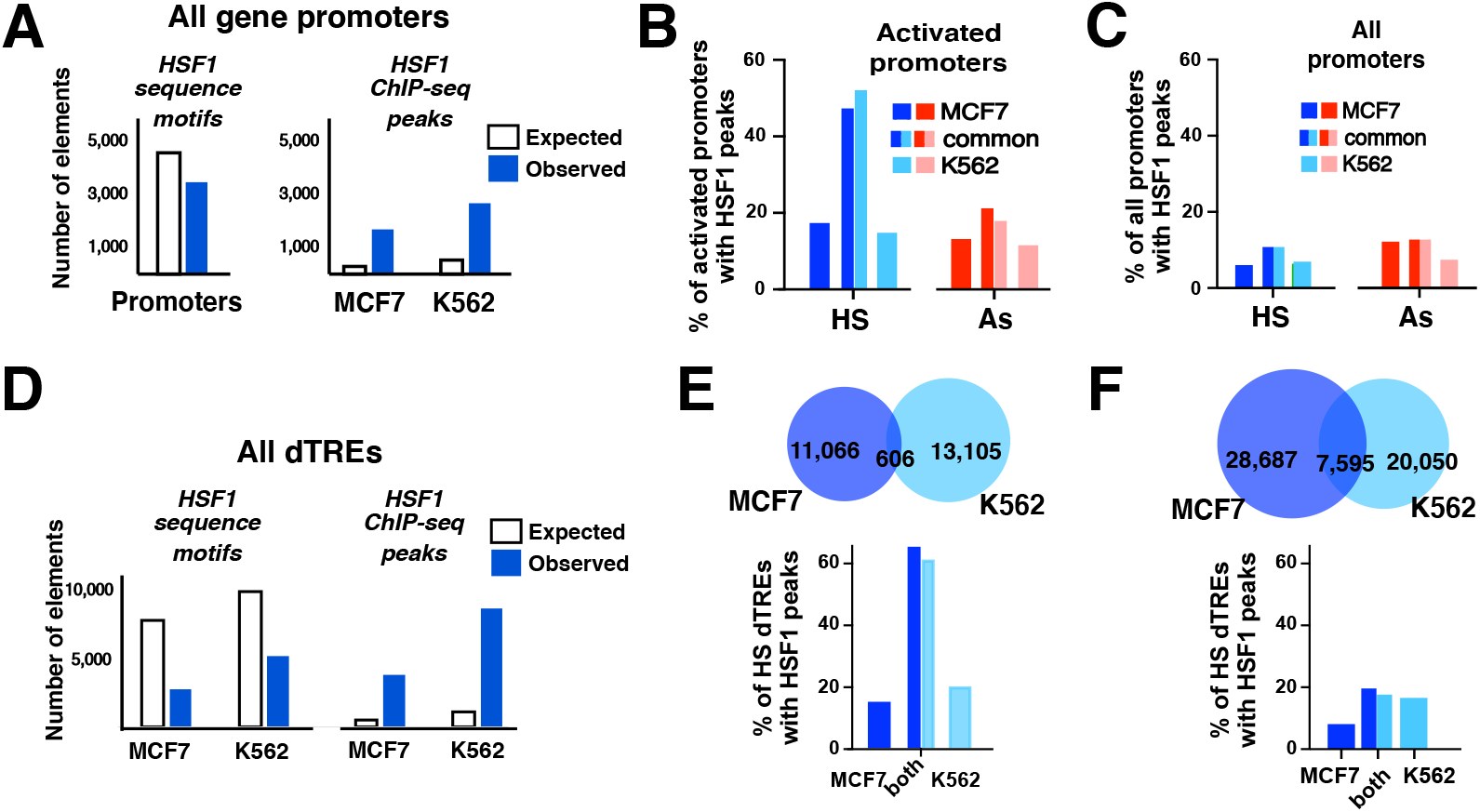
Genome-wide connection between HSF1 binding and function. **a. Left** panel. Bar graphs showing the numbers of actual (blue) HSE motifs at promoter regions (+/-1000nt from the TSSs) based on homer’s findmotif output for hg19 genome or the number of motifs expected within +/-1000nt from all TSSs assuming even distribution of findmotif output of ~286K motifs across hg19 genome (white). Right panel: bar graphs showing the numbers of experimentally observed (blue) HSF1 ChIP-seq peaks or the numbers of HSF1 ChIP-seq peaks based on even distribution of the same number of peaks across the genome (white). **b.** Fractions of activated promoters in HS (blue) and As (red) in MCF7 (dark blue, dark red) and K562 (light blue, light red) cells. Approximately 40% (in HS) and 20% (in As) of commonly activated genes showed HSF1 at their promoters, while less than 20% genes activated exclusively in MCF7 or K562 did. **c.** Percentages of HSF1 peaks at all gene promoters in HS (blue) and As (red) in MCF7 (dark blue, dark red) and K562 (light blue, light red) cells, at promoters common to both or exclusive to each cell line. **d.** Bar graphs showing the numbers of expected at ransom (white) and observed (blue) HSE motifs (left panel), and HSF1 ChIP-seq peaks (right panel) at all dTRES (dTREs defined as a region +/-1000 from dTRE center) in MCF7 and K562 cells. **e.** HS-activated dTREs between K562 and MCF7 cells. Venn diagram showing their overlap between the two cell lines (top). The percentage of dTREs with HSF1 peaks for dTREs common between the cell lines and exclusive dTREs for each cell line (bottom). **f.** Same as in (e) except all dTREs are shown.

Outside of promoters, HS was shown to involve intergenic sites previously referred-to as distant Transcription Regulatory Elements (dTREs) (30). We identified dTREs in our HS PRO-seq datasets based on nascent RNA signatures (Supplementary File 6). We do not refer to these elements as enhancers, although the identified dTREs overlapped with about 50% of enhancers previously described in either cell line (34). The proportion of dTREs with HSF1 peaks was also higher among dTREs unique to HS compared to NHS conditions (Supplementary Fig. 13). Examining DNA sequences, we noted an enrichment of HSF1 motifs around HS-induced dTREs (Supplementary Fig. 13c). However, the density of HSF1 peaks at dTREs was higher than the genomic density of HRE sequence motifs (Fig. 5d), indicating that, similar to promoters, HSF1 is mainly recruited to dTRE sites by means other than the HRE sequences. Like promoters, most dTREs were activated in the absence of HSF1 binding (Supplementary Fig. 13). A group of HS-activated dTREs common between the two cell lines showed the enrichment of HSF1 peaks that was at least as high as that for promoters of commonly activated genes (Fig. 5b, e). Gene Ontology (GO) analysis for genes associated with these dTREs that were previously annotated as enhancers (34) did not show statistically significant gene categories, but did include HS-induced heat shock protein *HSPA90* and *HSPA8* genes. However, this group of conserved dTREs was small (Fig. 5e, Supplementary Fig. 14) and did not sway the overall disconnect between HSF1 binding and transcription (Fig. 5f). The proportion of dTREs containing HSF1 peaks was similar to that at promoters (Fig. 5c, f), reinforcing a modest connection between HSF1 binding and transcription activation as a genome-wide property of HSR.

### HSR involves amplification of basal HSF1 binding

To understand how the genome-wide HSF1 binding may be established from the ground state, we compared HSF1 signal during HSR and at non-heat shock (NHS) conditions, considering two possibilities. First, HSF1 may occupy distinct sites before and during HSR. A vast majority of HSF1 peaks detected in NHS datasets were in common with HS samples (Fig. 6a, b). A small number of peaks unique to NHS (Fig. 6a, Supplementary File 7) were found mostly at promoters (42 out of 59 NHS-exclusive peaks in MCF7 cells compared to HS). The corresponding genes were overrepresented among metabolism-related GO categories and were not activated in HS (Supplementary File 7). Since NHS-exclusive binding involves only a small number of HSF1 peaks, we additionally searched NHS datasets for high intensity HSF1 signal regardless of specificity. A group of ~60 peaks around the centromeric regions of chromosomes showed high HSF1 signal (Fig. 6b, Supplementary Fig. 15). These regions showed pileups of Pol II and H3K4me3 signal as well (Supplementary Fig. 15) and may reflect the binding of proteins unrelated to their primary functions (35). This HSF1 signal did not change in HS (Fig. 6b) and, moreover, aligned along the slope of 1 in scatterplots, validating, if fortuitously, sequencing depth-based normalization of our ChIP-seq datasets. NHS-specific events thus constitute at most a tiny proportion of HSF1 binding.

**Fig. 6.**
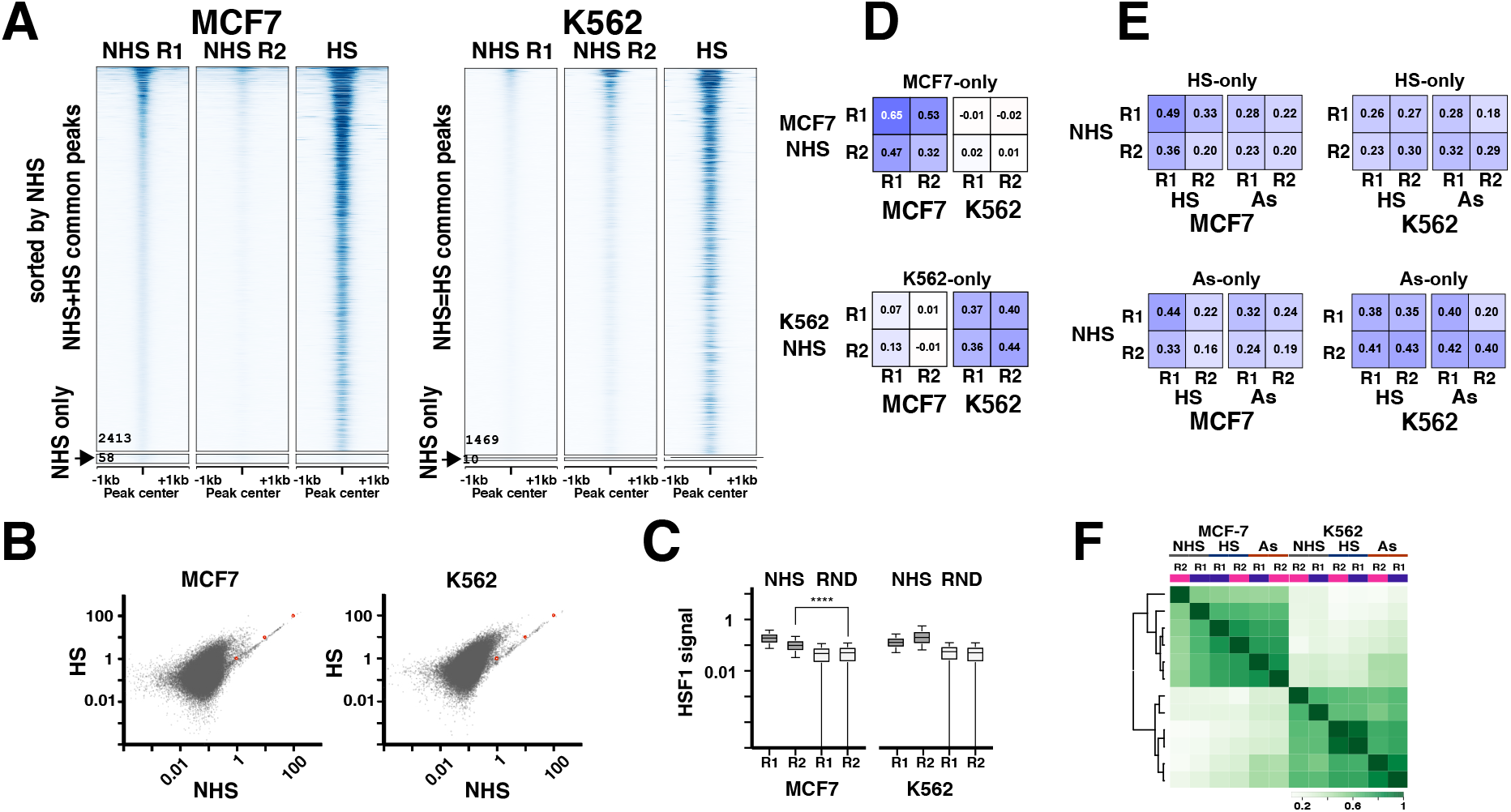
Comparison of basal and activated HSF1 binding. **a.** Heatmaps showing HSF1 peaks identified in NHS cells including peaks found in common with HS or As and a small number of peaks found exclusively in NHS cells (indicated by an arrow). Heatmaps are shown for each NHS replicate. **b.** Scatterplots of NHS and HS cells. Red dots indicate the slope of 1 following sequencing depth normalization. **c.** Averaged signal for NHS datasets from the locations of HS-only peaks not found in NHS cells compared to the signal for the same datasets from randomly selected genomic locations of the same average size and number (15,698 for MCF7 and 30,360 for K562). Random locations were defined using bedtools random with the average size of HSF1 peaks and filtered against promoters, dTREs, and HSF1 peak locations. The values for HS-only and random peaks are significantly different for each replicate (Mann-Whitney test, p<0.0001). **d.** Spearman correlation for signal in cell line-exclusive peak locations between NHS and HS cells for each cell line, shown for MCF7-only peaks (top) and K562-only peaks (bottom). **e.** Spearman correlation as in (d) except that HS and As-exclusive peak locations are shown for each cell line. **f.** Unbiased clustering of samples based on all HSF1 peaks across all datasets per replicate.

Another possibility is that HSF1 may bind to the same sites before and after HSR. In this case, low-intensity HSF1 signal should be evident in NHS cells at the locations of HSF1 peaks that would be newly acquired in HSR. The average HSF1 signal in NHS cells at HS peak locations, including HS-only peaks, was above that at randomly selected background regions (Fig. 6c). Unlike the peak regions, HSF1 signal at these background regions did not increase in HS (Fig. 6c, Supplementary Fig. 16). Amplification of low-level binding, therefore, appears to be the primary mode of HSR. This amplification can be either uniform for all sites, or selective. To address this, we compared HSF1 signal in NHS cells at the locations of HSR peaks exclusive to each cell line or treatment. NHS signal at cell line-exclusive peaks positively correlated with HSR signal in the same, but not the other cell line (Fig. 6d), indicating cell line specificity of basal HSF1 binding. Within a cell line, however, NHS signal at condition-specific peaks correlated with HSR signal in either treatment (Fig. 6e). These data indicate no preference of basal HSF1 binding prior to exposure to a stimulus and thus some selectivity of HSF1 signal amplification. However, these differences were not sufficient to affect the overall clustering of NHS samples by the cell line rather than treatment (Fig. 6f), indicating that HSF1 signal amplification is largely uniform. Furthermore, that the NHS samples retained cell line separation despite the low signal indicates that HSF1 binding patterns retain their identity across a broad dynamic range.

### HSF1 further opens remote inactive sites

Because chromatin defines how transcription factors interact with the genome (36, 37), we asked how HSF1 binding might relate to chromatin accessibility and, to that end, compared ATAC and HSF1 enrichment before and during HSR. A positive control *HSPH1* gene showed an HSR-dependent increase in ATAC signal along the gene body, consistent with highly activated transcription (Fig. 7a; Fig. 1a). However, HSR induced no drastic ATAC changes globally, as ATAC signal at the locations of peaks uniquely called in any one dataset, including NHS, was visually apparent in all of them (Fig. 7b). HSF1 was enriched in HS-only, but not As-only ATAC peaks (Fig. 7c). Statistical comparison revealed fewer changed ATAC peaks (p adj<0.05) and showed an overall increase in HS and decrease in As compared to NHS samples, with HSF1 in either treatment associated with an increase in ATAC signal (Supplementary Fig. 17). Over 40% of ATAC peaks that were statistically changed in HSR overlapped with dTREs (92 out of 216 in HS and 1,105 out of 2,613 in As) compared to only 3% overlapping with promoters (8 out of 216 in HS and 70 out of 2,613 in As). In NHS cells, ATAC peaks at promoters showed higher signal than at intergenic sites (p<0.0001) (Fig. 7d). In HS, an increase of ATAC signal was observed at intergenic sites (Fig. 7d, right), but not at promoters (Fig. 7d, left). The intergenic ATAC signal in HS was consistently higher at peaks containing HSF1 compared to peaks without HSF1 binding (p<0.0001) (Fig. 7d, right, blue bars). Less significant differences in ATAC signal were found between these peaks in NHS cells (Fig. 7d, right, grey bars). This resulted in a modestly yet consistently higher fold-increase of ATAC signal at sites containing HSF1 peaks (Fig. 7d, right). We conclude that HSF1 binding contributes to further chromatin opening at intergenic regions.

**Fig. 7.**
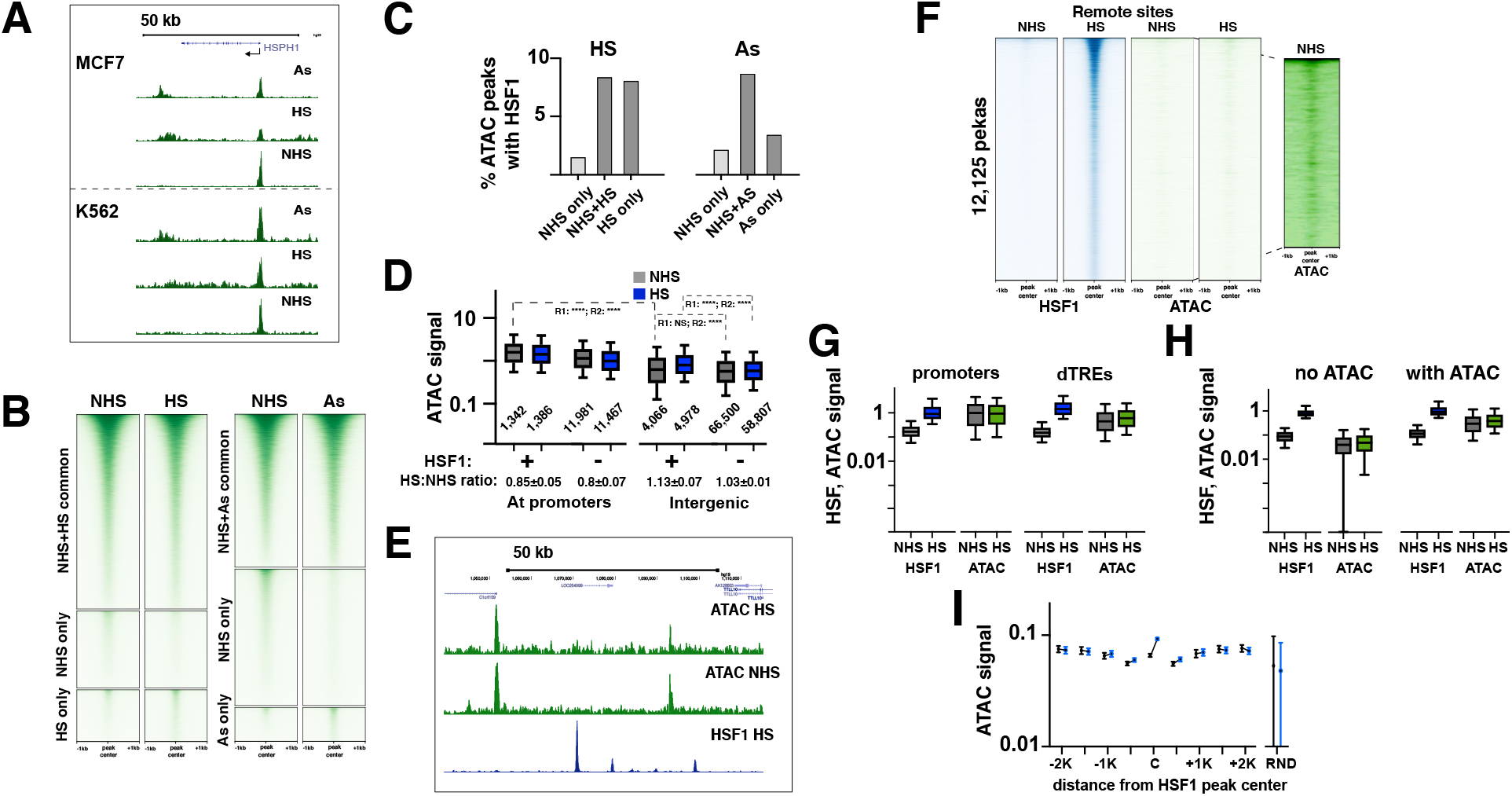
HSF1 favors locally open chromatin. **a.** UCSC browser ATAC tracks of *HSPH1* gene region for indicated conditions. **b.** Heatmaps showing common and condition-specific ATAC peaks in MCF7 cells between NHS and HS treatment (left) and NHS and As treatment (right). **c.** Percentage of ATAC peaks with MCF7 HSF1 peaks among ATAC peaks for common and condition-unique categories defined in (b). **d.** ATAC signal for ATAC overlap with promoters and presence (+) or absence (-) of overlapping HSF1 peaks. The numbers of ATAC peaks for each category are indicated below each boxplot. Also shown are the ratios of HS:NHS signal for each peak category based on two independent biological replicates (mean and range). Mann-Whitney test is shown for remote peaks for each replicate. **e.** A UCSC browser shot of a representative ~80-kb region showing ATAC and HSP1 signal in MCF7 cells. **f.** The heatmaps show HSF1 and ATAC signal in the regions of remote HSF1 peak sites outside of promoters, dTREs and annotated enhancers, for MCF7 cells at NHS and HS conditions. The inset shows a contrast-adjusted signal for NHS ATAC heatmap indicating pre-existing ATAC signal. **g.** HSF1 and ATAC signal for promoters (1,695) and dTREs (3,900) overlapping with HSF1 peaks, for NHS and HS datapoints. **h.** HSF1 and ATAC signal for remote genomic HSF1 peak regions in (f) that overlap (2,132) or do not overlap (9,994) with ATAC peaks. Signal in (g) and (h) is shown in the same scale for each dataset. The numerical similarity of scales between HSF1 and ATAC datasets is coincidental. **i.** ATAC signal at the locations of 9,994 distant MCF7 HSF1 peaks acquired in HS, shown as the mean with 95% confidence interval. The averaged ATAC signal centered on HSF1 peaks is shown for 500-nt bins around the peak center (“C”) with the offset from the center indicated. NHS ATAC signal is shown in black and HS in blue.

To identify the preference of HSF1 for pre-existing open versus closed chromatin, we examined the ATAC signal across the HSF1 peak sites. Visually, we noted a widespread lack of the overlap between HSF1 and ATAC signal pileups (Fig. 7e). Only about 1/3^rd^ of all HSF1 peaks in MCF7 HS datasets overlapped with NHS ATAC peaks, with 2/3^rd^ of this overlap accounted for by promoters and dTREs (Supplementary Fig. 18). As expected, HSF1 peaks found at promoters or dTREs bound to sites that were open before HS (Supplementary Fig. 19). Outside of these elements, however, HSF1 appeared to bind to closed chromatin (Fig. 7f). Indeed, most of these remote HSF1 peak sites did not contain ATAC peaks and showed no active histone marks (Supplementary Fig. 20). However, careful examination revealed low-level yet widespread pre-existing ATAC signal in NHS cells around the locations of HSF1 peaks that would be acquired in HS (Fig. 7f, inset). These observations are consistent with HSF1 favoring pre-existing sites. To gain insight into HSF1 binding, we compared HSF1 binding intensity to genomic sites with various degrees of ATAC signal. Whereas ATAC signal was higher at promoters than dTREs, consistent with promoters being more open, HSF1 signal was higher at dTREs (p<0.0001) (Fig. 7g). Likewise, comparing open and closed remote HSF1 binding sites distinguished by ATAC, HSF1 signal was disproportionally higher at more closed sites (Fig. 7g, h). HSF1 binding, therefore, cannot be fully explained by the magnitude of the pre-existing ATAC signal. We noted a higher prevalence of HRE motifs among HSF1 peaks at more closed remote regions (Supplementary Fig. 21). The binding of HSF1 to more closed chromatin may thus be more reliant on direct DNA sequence-specific interactions.

Lastly, we examined remote HSF1 peaks for changes in chromatin accessibility that might potentially occur in HS. These sites showed the ATAC signal increase in HS at the exact locations of HSF1 peaks, but not in the immediate surrounding regions (Fig. 7i). This increase in ATAC signal in HS was modest as it did not result in ATAC peaks (Fig. 7f, i). Nevertheless, co-occurrence of this increase with HSF1 binding locations lends support to HSF1 binding independent of the ChIP method. These data also suggest that HSR can trigger changes in remote genomic regions.

## DISCUSSION

Inducing distant cell lines with two stimuli involving a common transcription factor allowed us to define the cell type specificity and robustness for its genome-wide binding as well as connection to transcription. In contrast to *Drosophila,* in which transcription in heat shock is globally inhibited in favor of few highly activated loci (1, 6, 30, 38), mammalian transcription continues across the genome. This as well as the numbers of genes transcriptionally activated during HSR are in line with responses to other environmental and physiological signals (23, 39–41). Whether induced by heat or chemically, mammalian HSR is not quantitatively different from responses to other signals. TF binding patterns reflecting the activity of signaling pathways (23, 40, 41) may serve as readouts of cellular identity, building, for example, on previously noted cancer-specific HSF1 programs (3).

That genome-wide HSF1 patterns show cell line specificity even prior to activation argues against distinct basal-specific HSF1 programs and, we suggest, in general against the diversity of TF binding patterns in a given cell type. It is conceivable that low-level basal HSF1 binding (Fig. 5) may have resulted from inadvertent activation of HSR during routine cell culturing. However, our basal readouts are in line with those reported previously (6). This binding also indicates that HSR is not an all-or-nothing response, but is commensurate with the stimulus intensity. Retention of genome-wide patterns at different magnitudes of activation suggests that HSF1 is not hoarded at specific high-affinity sites at the expense of others, but is distributed linearly across the genome based on the amount of active HSF1 in the cell. Mechanistically, this is consistent with rapid sampling of available sites by individual HSF1 molecules (42, 43). This also in agreement with previous findings that HS causes no major changes in the nuclear 3D architecture (30, 44). Our ATAC data show further that the architecture is preserved in HSR down to individual loci. Even though how exactly HSF1 binding patterns echo the nucleus remains to be defined, the binding of HSF1 to existing functional elements such as promoters or enhancers appears to be driven to a higher extent by co-factors or favorable structure whereas the binding to inactive remote sites outside of functional elements may rely more on the DNA sequence.

Transcription factor binding versus function is a fundamental question that is yet to be answered in any system. HSF1 binding during HSR is decoupled from transcription activation and, beyond few well known loci, shows little conservation between distant cell lines. This disconnect is overwhelmingly due to variability of transcriptional outcomes rather than HSF1 binding. Some of this disconnect is due to HSF1 action at promoters while binding at distant enhancers (45). However, the degree of association of HSF1 binding and nearby transcription activation is similar at and outside of promoters, suggesting that the effect of HSF1 on transcription is mainly local. This may reflect combinatory regulation wherein the transcription output is defined by context-specific combinations of individually stable transcription factor patterns. Apart from HSF1, response to heat has been shown to involve transcription factors including HSF2, SRF, CTCF, ER-α among others (5–7, 46–48), whereas As may also involve oxidative stress components (49, 50). Notably, recent work demonstrated that deletion off HSF2 does not dramatically alter HSF1 binding (7).

Another question is about functionality of low-affinity TF binding sites that are progressively identified in omics studies. At least some of low-affinity TF binding sites likely are functional (51), with cooperativity among individual binding events driving the binding across the genome including enhancers (52, 53). The HS-dependent increase in ATAC signal at remote HSF1 binding indicates engagement of chromatin independent of transcription. TF binding may nucleate the opening of closed genomic regions, whether immediately, over time or with repeated stimulation. HSR is often ectopically activated in cancers: arsenic is a transforming agent that in addition to mutations can induce epigenetic changes (54), and heat has been recently shown to induce transcriptional memory (45). Changes prompted by stimuli should depend on transcriptional responses to activate transcription factor binding to DNA, but not necessarily transcription (55).

## METHODS

### Cell culture and treatments

All cells were purchased from the American Type Culture Collection and used within the first 15 passages. Cells were grown in 15cm dishes as before (56) and growth media was replaced with fresh media 24 hours before treatments (30, 57). Heat-Shock treatment was started by placing a dish onto a heated water surface for 1 minute to raise the temperature quickly and continued in a 42°C 5% CO_2_ incubator for the remaining time. Arsenic treatment was performed by the addition of 500 μM Sodium meta-arsenite (Sigma) followed by incubation at the ambient 37°C with 5% CO_2_ (24).

### Western blotting

Cells were scraped (MCF-7) or collected from suspension (K562), washed with cold 1x Phosphate-Buffered Saline (PBS), resuspended in lysis buffer (8M Urea, 1% SDS, 126mM Tris pH 6.8), and protein concentration was measured on Qubit fluorometer using protein assay (Invitrogen). Approximately 30μg of total protein was resolved on a 10% SDS-PAGE gel and transferred onto a PVDF membrane. Nonspecific binding was blocked by 5% nonfat milk diluted in 1x TBS with 0.1% Tween-20 (TBS-T) followed by primary antibody incubation overnight and three TBS-T washes. Blots were developed using horseradish peroxidase-conjugated secondary antibody (GE) and imaged on a Li-COR Odyssey Fc imager. Western blotting was performed against HSF1 (Enzo # ADI-SPA-901-D) or [pSer^326^]-HSF1 (Enzo # ADI-SPA-902-D) antibodies and GAPDH (Millipore # ABS16) as control. Band sizes were verified using Bio-Rad Precision Plus dual color protein ladder.

### Reverse transcription and quantitative PCR

Approximately 500 ng of total RNA was used to synthesize cDNA using random hexamers and SSRT III reverse transcriptase (Life Technologies). Quantitative PCR data were normalized against GAPDH gene transcripts and shown as fold change from at least three independent biological replicates. Error bars represent standard error of the mean.

### ChIP-sequencing

Approximately 2×10^7^ cells were crosslinked with 1% ethanol-free formaldehyde (Thermo) in serum-free DMEM/F12 media at 25°C for 10min, quenched with 125mM Glycine, and lysed in buffer (1% Sodium-Dodecyl Sulfate (SDS), 50mM Tris (pH=8), 10mM EDTA, 1X PMSF and protease inhibitor cocktail) followed by disruption in the Covaris S2 sonicator for 6 minutes (with peak power = 140, Duty factor = 5, Cycles/burst = 200). Protein A and Protein G Dynabeads (Invitrogen) premixed at 1:1 ratio were used for pre-clearing for 1 hour at 4°C. One percent of each precleared sample was taken as input, with at least two inputs taken for each sample and averaged for ChIP percent input calculation. Samples were incubated with 5μg of primary antibodies in 2 ml of dilution buffer (1% Triton X-100, 2mM EDTA, 20mM Tris pH=8.0, 150mM NaCl, 1X Phenylmethylsulfonyl fluoride (PMSF) and cOmplete protease inhibitor cocktail (Sigma)) with slow rotation overnight at 4°C followed by washing as before (56, 58). Four percent of DNA after ChIP was taken for validation by qPCR using positive and negative control primers (Supplementary File 8) to calculate the percent input as well as positive to negative control signal ratios for quality control of ChIP. DNA libraries were prepared from validated ChIP material using NEBNext Ultra II DNA library kit (New England Biolabs) protocol without size selection. Final PCR amplification of cDNA templates was done using TruSeq Illumina Small RNA PCR primers, 250 μM dNTP mix, 1x HF buffer, 1M Betaine, and Phusion DNA polymerase (NEB). Reactions were supplemented with 1xEvaGreen dye (Biotium) to monitor amplification and were manually terminated within the linear range of amplification, which was normally reached after 10 to 13 PCR cycles. Libraries were additionally validated by qPCR with the same primers (Supplementary File 8) by calculating positive to negative control signal ratios prior to sequencing (58). ChIP-sequencing was performed with anti Rpb1 NTD (D8L4Y) (CST #14958), HSF1 (Enzo #ADI-SPA-901-D) or H3K4me3 (Abcam #ab8580). Where indicated, experiments were done using anti HSF1 antibody (SCBT #sc-9144, no longer available).

### PRO-sequencing

Precision Run-On sequencing (PRO-seq) was performed as previously described (59). Briefly, nuclei were extracted from ~2×10^7^ cells and run-on reactions were carried on at 37°C for 3 minutes using 3μl of each 11-biotin-labelled ribonucleotide stocks (Perkin Elmer). Following real time PCR amplification in the presence of EvaGreen dye as above, libraries were run on a 6% TBE gel (Novex) with 1X Tris-Borate-EDTA (TBE) buffer. The gel was stained with ethidium bromide, visualized with a 312nm UV transilluminator, and areas between ~125-300bp DNA sizes were cut out and extracted using crush and soak method as done before (60).

### ATAC-sequencing

The Assay for Transposase Accessible Chromatin with high-throughput sequencing (ATAC-seq) was done using the original protocol and buffer (25). Incubation with Tn5 transposase was done in a shaking heating block at 37°C and 500 RPM for 30 minutes followed by real-time PCR amplification with Nextera dual index primers and NEBNext High-Fidelity 2x PCR mix with added Evagreen dye to avoid overamplification.

### ChIP-seq data analysis

Sequencing was done at the UND Genomics Core or commercially to the depth indicated in Supplementary File 1. H3K27Ac and H3K4me1 datasets were from a public dataset under GEO accession number GSE85158 (61). Illumina adapters were removed from raw files with trimmomatic using paired end mode and keepBothReads set as ‘true’ and aligned to hg19 genome in paired end mode with hisat2 using --no-spliced-alignment and otherwise default parameters (62). BigWig files were made using bamCoverage (63). For all ChIP-seq data except HSF1 and for HSF1 scatterplots, read counts were normalized to the sequencing depth and signal was calculated from the bigwig files with multiBigwigSummary with sequencing depth coverage normalization. For HSF1 peak calling, alignments were filtered against blacklisted genomic regions that included ENCODE ENCSR636HFF excluded list regions (64), regions known to be absent (zero copy number) in either MCF7 or K562 cell lines from Cancer Cell Line Encyclopedia (65), and UCSC blacklisted human genome regions (66). After removing PCR duplicates with MarkDuplicates (Picard), the resultant bam files were subset using samtools view to the sample with the lowest coverage based on the numbers of primary reads (-F 260 option) in the sample, and peaks were called using macs2 (67) with FDR cutoff 0.01. UCSC genome browser tracks were generated using sequencing depth-normalized bigwig files, with each type of a track shown in the same numerical Y scale for all samples. Individual peak list replicates were merged by combining peaks present in both replicates (K562 cells) or either replicate (MCF7 cells). Differential signal was calculated using DiffBind (68). PCA plots were generated using affinity values default function for PCA in DiffBind. The 23K (n = 23,698) gene list and exact genes and TSS definitions were as defined previously (30), with promoter regions defined as TSS+/-1000nt. Motif search for HSF1 was done with HOMER (22). Venn diagrams were drawn using R venneuler package.

### PRO-seq and ATAC-seq data analysis

Illumina small RNA TruSeq adapters were trimmed and reads shorter than 15 nt were discarded. Trimmed reads were used to remove ribosomal RNAs and remaining reads were aligned to hg19 using hisat2 using no-splicing option. Individual samples were normalized using 3’-ends of long genes where the elongating Pol II could not have reached within the timing of treatment (59). Differentially expressed genes were defined using DESeq2 (p adj<0.05). Identification of dTREs was done using the remote site (30) with default parameters for individual replicates, retaining dTREs present in both biological replicates for MCF7 cells and one replicate for K562 cells. HS-specific dTREs were defined as present only in HS and not in NHS cells. For ATAC-seq, adapter-trimmed reads were aligned to hg19 genome as above, followed by converting to BigWig format using bamCoverage with the default bin size of 50 bp. The signal was normalized using counts per million (CPM) and duplicate reads were removed using MarkDuplicates. Heatmaps were generated using deepTools plotHeatmap.

### Statistics

Peaks were called in each replicate using macs2 with q-value cut-off of 0.01. Differentially expressed genes were defined using DeSeq2 based on p_adj 0.05 using on two replicates unless indicated otherwise. Differential ChIP-seq and ATAC peaks were defined based on DiffBind p_adj 0.05 using otherwise default settings. Quantitative PCR was based on three or more independent biological replicates for RT-qPCR and ChIP-qPCR and plotted as SEM+/-SD. Omics comparisons were plotted with error bars showing 10-90 percentiles. Comparisons of HSF1 and PRO-seq signal were done using nonparametric Mann-Whitney test calculating two-tailed p-values.

## Availability of data and materials

Original datasets are available in the Gene Expression Omnibus repository under <u>GSE209687</u> accession number. Public datasets analyzed during the current study are under accession number <u>GSE85158</u>.

## Authors’ contributions

S.G.D. – designed the study, performed experiments to generate most datasets, analyzed and interpreted data, and wrote the manuscript. B.D.K. – analyzed HSF1 ChIP-seq and ATAC data for differential signal and interpreted the analyses. B.L. – performed Western blots. D.Pa., D.Pe., M.V. – analyzed PRO-seq data. A.T. – generated PCA plots for ChIP-seq data. N.K.K.K. – prepared cells for PRO-seq experiments. A.A. – performed initial HSF1 ChIP-seq experiments with SCBT antibody. N.S. – designed the study, analyzed data and wrote the manuscript.

## Acknowledgements

We thank Archana Dhasarathy, Motoki Takaku, Min Wu, Benjamin Roche and Paul Wade for critical reading of the manuscript, members of Dhasarathy and Takaku labs for productive discussions throughout, as well as Sara Apostal and Mike Hill for handling omics libraries and data. We also thank the UND Genomics Core for prompt service. The work was supported by the National Science Foundation CAREER award 1750379 to NS and by the National Institute of General Medical Sciences of the National Institutes of Health under Award Numbers U54GM128729 and 2P20GM104360. The funding bodies had no role in the design of the study, collection, analysis, and interpretation of data or in writing the manuscript.

## SUPPLEMENTARY INFORMATION

**Supplementary Figures** 1-21

**Supplementary_file_1:** Excel (xlsx), Datasets used in this study including alignment statistics.

**Supplementary_file_2:** Excel (xlsx), Locations of HSF1 peaks for each condition including chromosome, peak start, peak end. All coordinates are based on hg19 genome annotation.

**Supplementary_file_3:** Excel (xlsx), Promoters of upregulated genes for each condition, including chromosome, interval start, interval end, gene symbol, strand. Intervals are defined as transcription start site +/-1000 nucleotides.

**Supplementary_file_4:** Excel (xlsx), Panther GO Ontology over-representation test for upregulated genes in MCF7 cells that are commonly activated in HS and As, unique to HS and unique to As.

**Supplementary_file_5:** Excel (xlsx), Panther GO Ontology over-representation test for HS upregulated genes in MCF7 and K562 cells including common between MCF7 cells, unique to MCF7 and unique to K562 cells.

**Supplementary_file_6:** Excel (xlsx), Coordinates of dTREs identified in this study in the cell line and condition indicated, including chromosome, dTRE start, dTRE end.

**Supplementary_file_7:** Excel (xlsx), Promoters associated with basal (NHS-only) peaks in MCF7 cells, including coordinates and names of the corresponding genes as well as GO Gene Ontology PANTHER overrepresentation test.

**Supplementary_file_8:** Excel (xlsx), Primers for qPCR used in this study.

