## Supplementary Figures for "Transcriptional Responses of Cancer Cells to Heat Shock-Inducing Stimuli Involve Amplification of Robust HSF1 Binding"

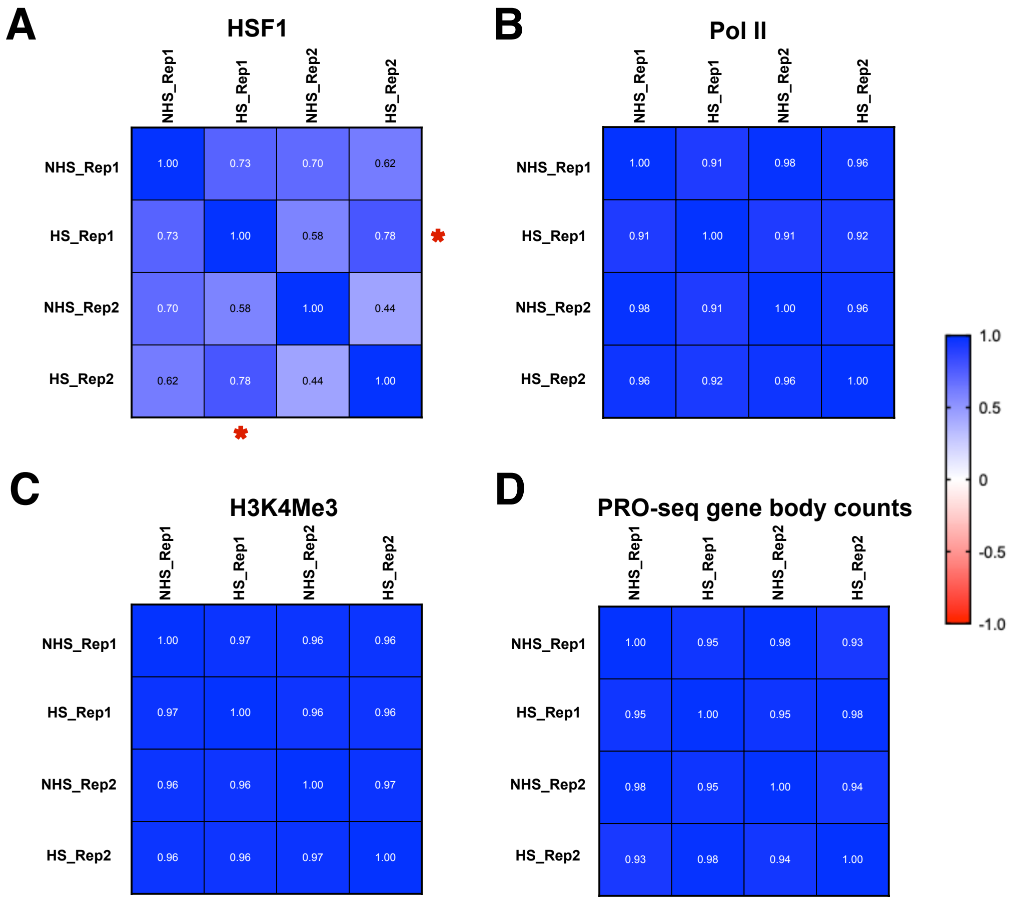


**Figure S1. Correlation among datasets.** Spearman correlation among datasets based on counts within peak regions for HSF1 ChIP-seq datasets, promoter regions (+/- 1000bp) for other ChIP-seq datasets, and gene body counts for PRO-seq datasets. Asterisks indicate correlation between MCF7 HS replicates mentioned in the main text.

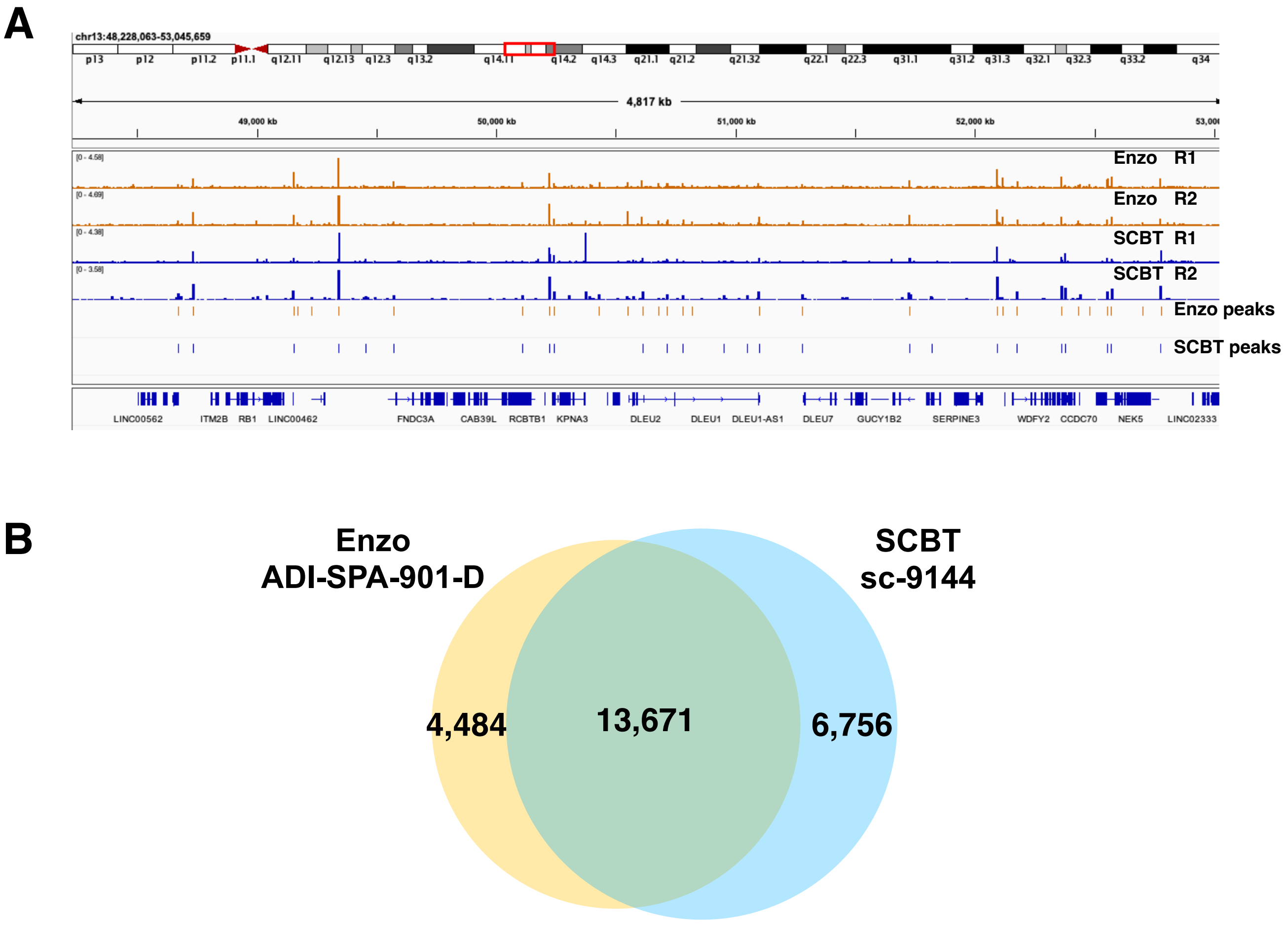


**Figure S2. Comparison of anti-HSF1 antibodies.** A. IGV browser ChIP-seq tracks with anti-HSF1 antibodies including monoclonal Enzo (orange, used throughout this work) and polyclonal SCBT as indicated (blue) showing a randomly selected ~5mB region of the genome, for each replicate. The polyclonal antibody is no longer available from any supplier. IGV browser tracks for HS peaks based on Enzo (used throughout this work) and SCBP anti pan-HSF1 antibodies. B. Venn diagram for MCF7 ChIP-seq peaks between the two anti-HSF1 antibodies.

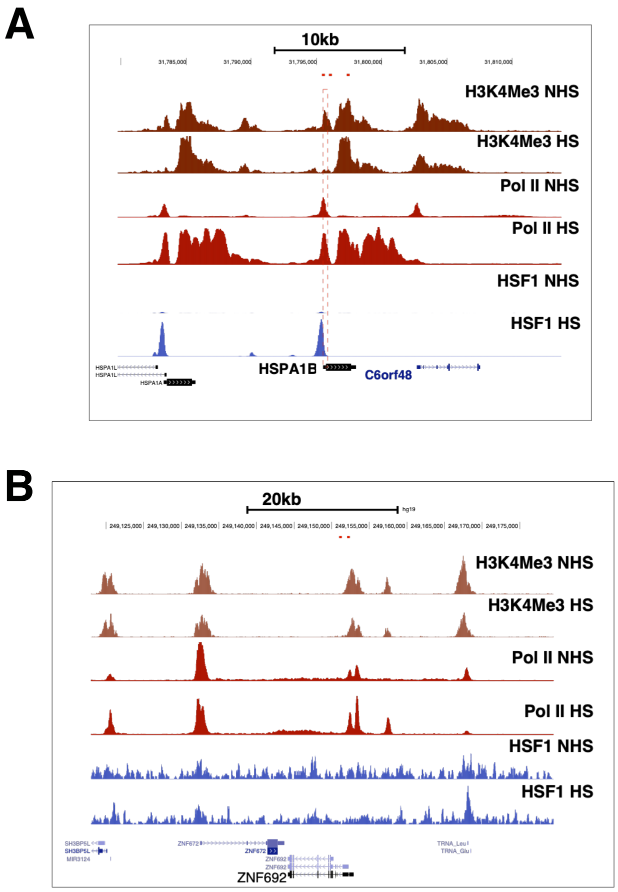


**Figure S3. H3K4Me3 tracks.** UCSC browser track showing a region around HSPA1B (HSP70) heat shock activated gene (A) and control gene that is not changed in HS (B). Sites of qPCR primers for these two genes are indicated above the tracks. H3K4Me3, Pol II and HSF1 tracks for the same regions are shown for MCF7 cells.

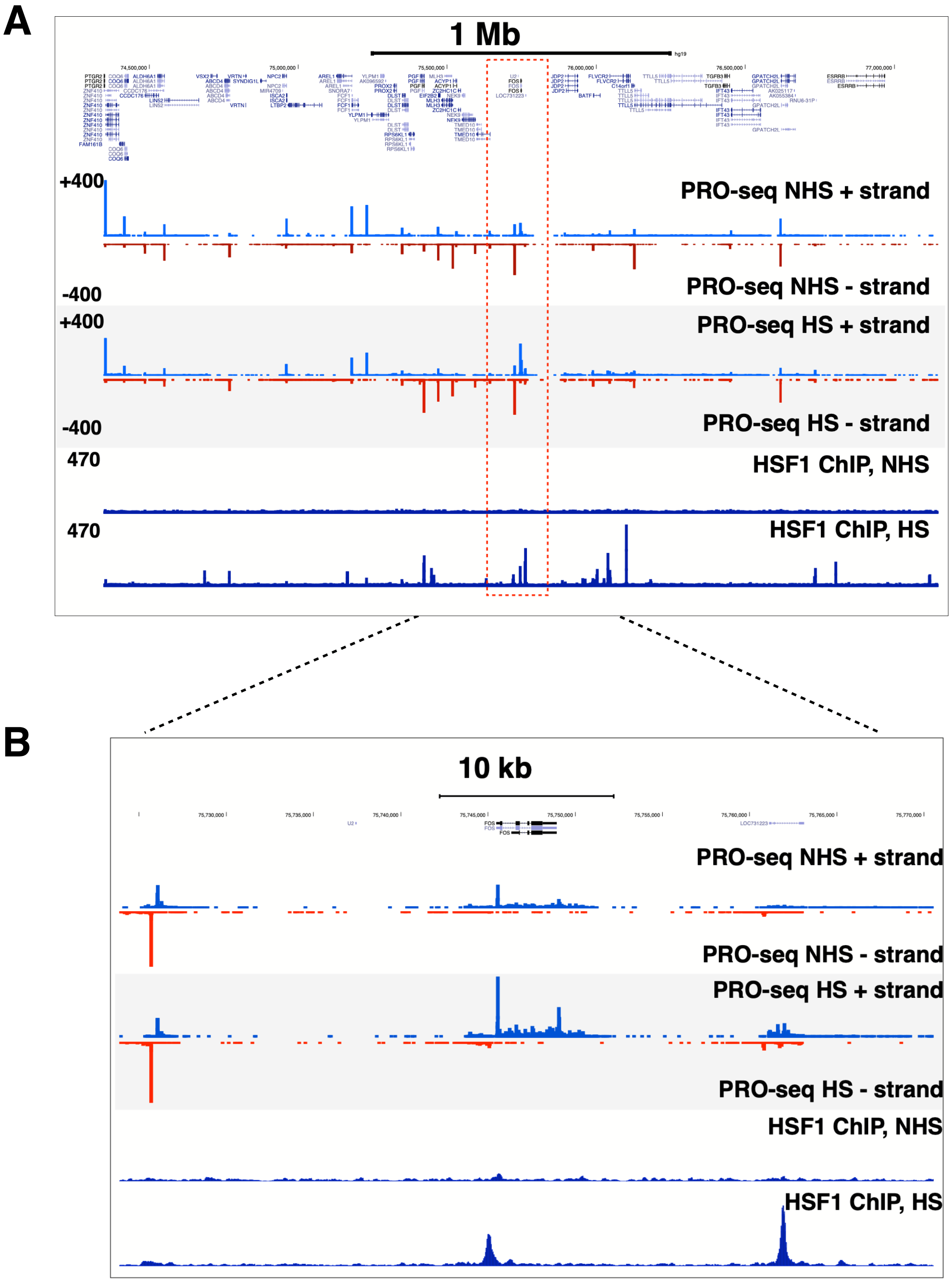


**Figure S4. PRO-seq tracks.** UCSC browser track showing PRO-seq and HSF1 ChIP-seq tracks for ~3mB region around a moderately HS-activated gene in MCF7 cells (FOS) (A) and zoomed in region around the same gene (B).

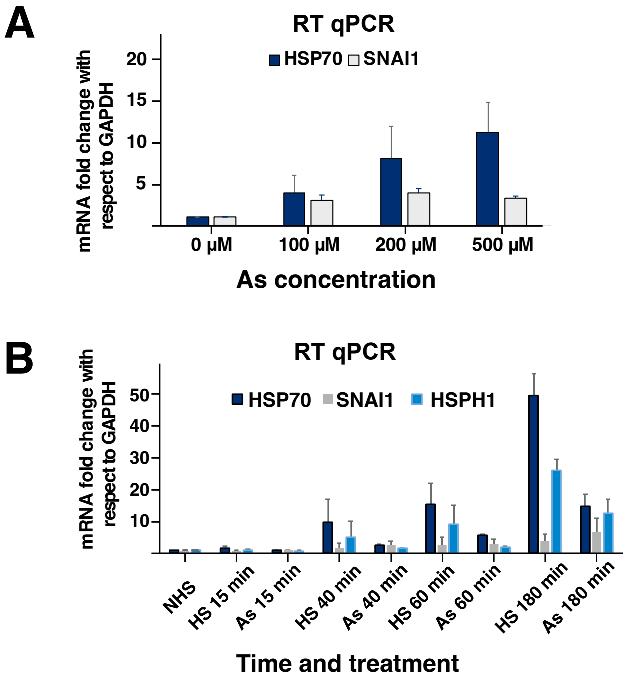


**Figure S5. Optimization of As-treatment.** (A) RT-qPCR showing titration of As concentration in MCF7 cells, with timing fixed at 60 minutes. (B). RT-qPCR showing titration of the time of As treatment. HS treatment was performed in parallel for the same time points and is shown next to As datapoints. Each experiment is based on at least three independent biological replicates.

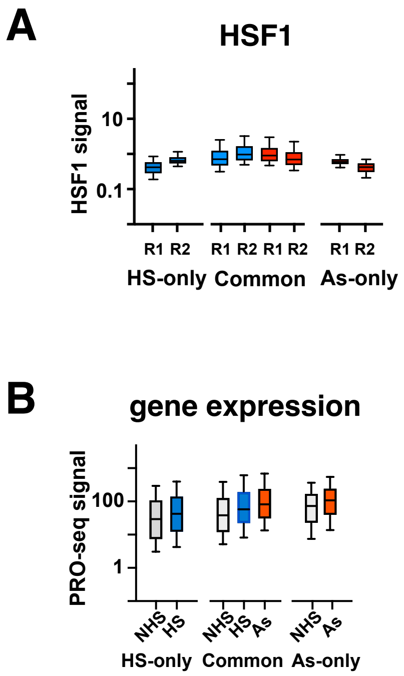


**Figure S6.** (A) HSF1 signal for MCF7 cells for peaks that are common for HS and As treatments, or unique to each treatment. HS signal is shown in blue and As in red. (B) PRO-seq gene body density signal for genes activated in both treatments or only in one treatment as indicated.

### Common HSF1 peaks

| Rank | Motif | Name | P-value | log P-pvalue | q-value (Benjamini) | # Target Sequences with Motif | % of Targets Sequences with Motif | # Background Sequences with Motif | % of Background Sequences with Motif |
|---|---|---|---|---|---|---|---|---|---|
| 1 | 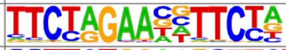 | HRE(HSF)/Striatum-HSF1-ChIP-Seq(GSE38000)/Homer | 1e-10422 | -2.400e+04 | 0.0000 | 9754.0 | 80.31% | 1750.2 | 4.66% |
| 2 | 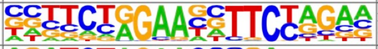 | HRE(HSF)/HepG2-HSF1-ChIP-Seq(GSE31477)/Homer | 1e-9084 | -2.092e+04 | 0.0000 | 8007.0 | 65.93% | 1057.1 | 2.82% |
| 3 | 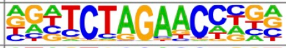 | ZBTB12(Zf)/HEK293-ZBTB12.GFP-ChIP-Seq(GSE58341)/Homer | 1e-931 | -2.144e+03 | 0.0000 | 3549.0 | 29.22% | 3233.3 | 8.61% |
| 4 | 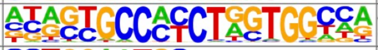 | CTCF(Zf)/CD4+-CTCF-ChIP-Seq(Barski_et_al.)/Homer | 1e-264 | -6.093e+02 | 0.0000 | 908.0 | 7.48% | 694.3 | 1.85% |
| 5 | 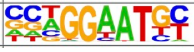 | TEAD4(TEA)/Tropoblast-Tead4-ChIP-Seq(GSE37350)/Homer | 1e-196 | -4.518e+02 | 0.0000 | 3692.0 | 30.40% | 7155.9 | 19.06% |

### HS-only HSF1 peaks

| Rank | Motif | Name | P-value | log P-pvalue | q-value (Benjamini) | # Target Sequences with Motif | % of Targets Sequences with Motif | # Background Sequences with Motif | % of Background Sequences with Motif |
|---|---|---|---|---|---|---|---|---|---|
| 1 | 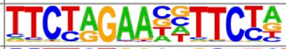 | HRE(HSF)/Striatum-HSF1-ChIP-Seq(GSE38000)/Homer | 1e-3595 | -8.280e+03 | 0.0000 | 3665.0 | 61.64% | 1587.0 | 3.64% |
| 2 | 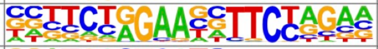 | HRE(HSF)/HepG2-HSF1-ChIP-Seq(GSE31477)/Homer | 1e-2559 | -5.894e+03 | 0.0000 | 2602.0 | 43.76% | 976.2 | 2.24% |
| 3 | 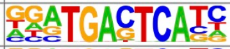 | Fra2(bZIP)/Striatum-Fra2-ChIP-Seq(GSE43429)/Homer | 1e-218 | -5.039e+02 | 0.0000 | 1144.0 | 19.24% | 2976.4 | 6.82% |
| 4 | 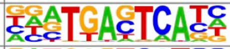 | Fra1(bZIP)/BT549-Fra1-ChIP-Seq(GSE46166)/Homer | 1e-209 | -4.833e+02 | 0.0000 | 1257.0 | 21.14% | 3574.4 | 8.19% |
| 5 | 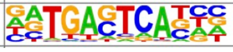 | Fosl2(bZIP)/3T3L1-Fosl2-ChIP-Seq(GSE56872)/Homer | 1e-200 | -4.623e+02 | 0.0000 | 853.0 | 14.35% | 1891.4 | 4.33% |

### As-only HSF1 peaks

| Rank | Motif | Name | P-value | log P-pvalue | q-value (Benjamini) | # Target Sequences with Motif | % of Targets Sequences with Motif | # Background Sequences with Motif | % of Background Sequences with Motif |
|---|---|---|---|---|---|---|---|---|---|
| 1 | 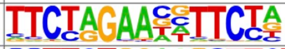 | HRE(HSF)/Striatum-HSF1-ChIP-Seq(GSE38000)/Homer | 1e-3480 | -8.014e+03 | 0.0000 | 3055.0 | 74.28% | 1539.7 | 3.41% |
| 2 | 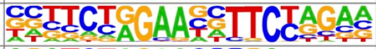 | HRE(HSF)/HepG2-HSF1-ChIP-Seq(GSE31477)/Homer | 1e-2980 | -6.864e+03 | 0.0000 | 2395.0 | 58.23% | 811.1 | 1.80% |
| 3 | 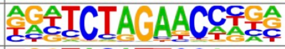 | ZBTB12(Zf)/HEK293-ZBTB12.GFP-ChIP-Seq(GSE58341)/Homer | 1e-298 | -6.878e+02 | 0.0000 | 944.0 | 22.95% | 2539.7 | 5.63% |
| 4 | 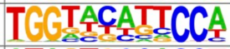 | PRDM10(Zf)/HEK293-PRDM10.eGFP-ChIP-Seq(Encode)/Homer | 1e-70 | -1.624e+02 | 0.0000 | 676.0 | 16.44% | 3582.0 | 7.94% |
| 5 | 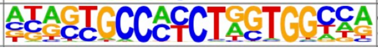 | CTCF(Zf)/CD4+-CTCF-ChIP-Seq(Barski_et_al.)/Homer | 1e-55 | -1.273e+02 | 0.0000 | 180.0 | 4.38% | 468.7 | 1.04% |

**Figure S7. Enrichment of transcription factor motifs for HSF1 peaks between HS and As treatments.** The top five transcription factor motif hits sorted by the p-value are shown as output of homer known motifs.

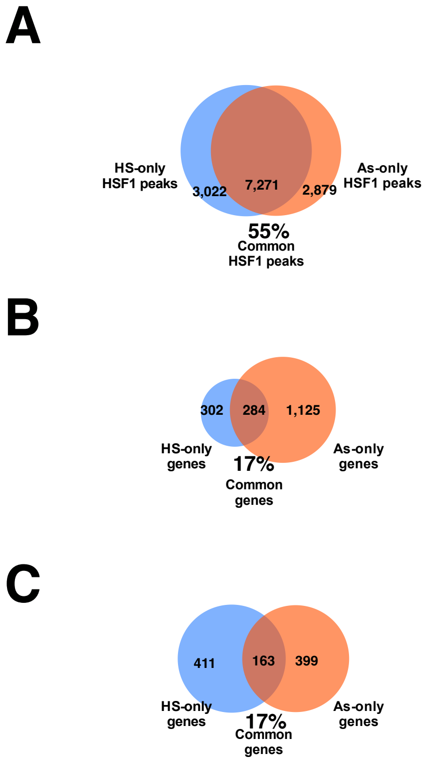


**Figure S8. Overlaps between HS and As treatments do not change with increased stringency.** (A) Venn diagram showing the overlap between HSF1 peaks observed in HS and As treatments in MCF7 cells showing only those peaks that were identified in both replicates. The percentage of common peaks among all peaks is shown underneath. (B). Venn diagram showing the overlap between activated genes as detected by PRO-seq using a minimum absolute gene expression (0.001 reads per million per kilobase) and a minimum fold change in at activated genes as 1.4. (C). Venn diagram of activated genes as in (B), but showing the same number of genes in HS and As conditions based on sorting the activated genes based on fold-change. The percentage of commonly activated genes among all genes is shown underneath.

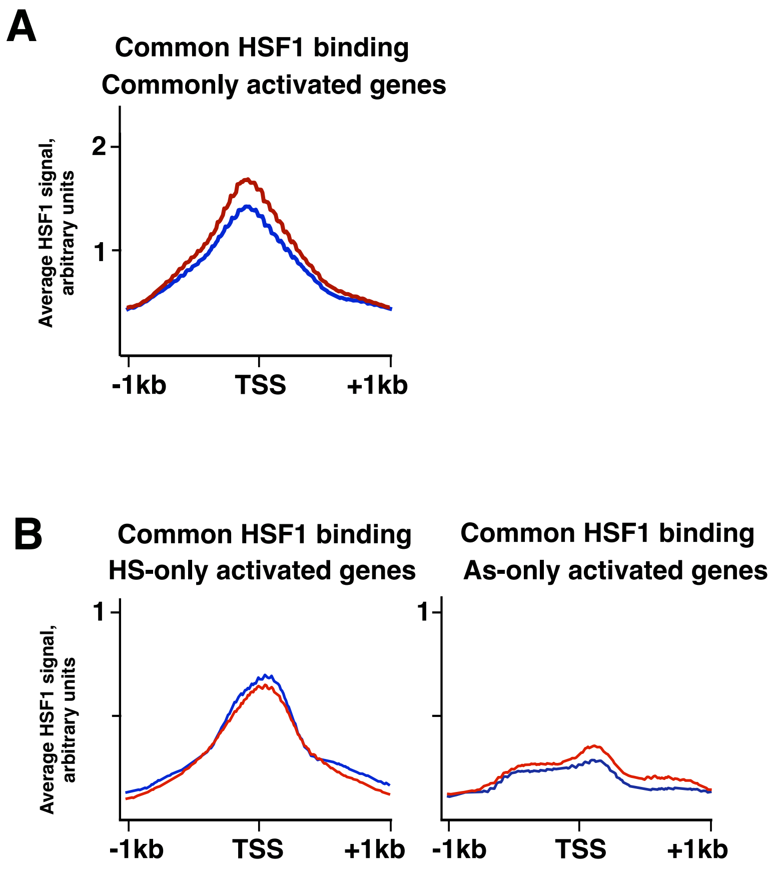


**Figure S9. HSF1 binds to the same sites between HS and As treatments.** Metaplot of HSF1 ChIP-seq profiles for promoters whose genes are activated in both treatments based on PRO-seq (A) or bound in both treatments, but bound in both treatments, but activated in only one (B). The Y-axis scale is similar for all plots.

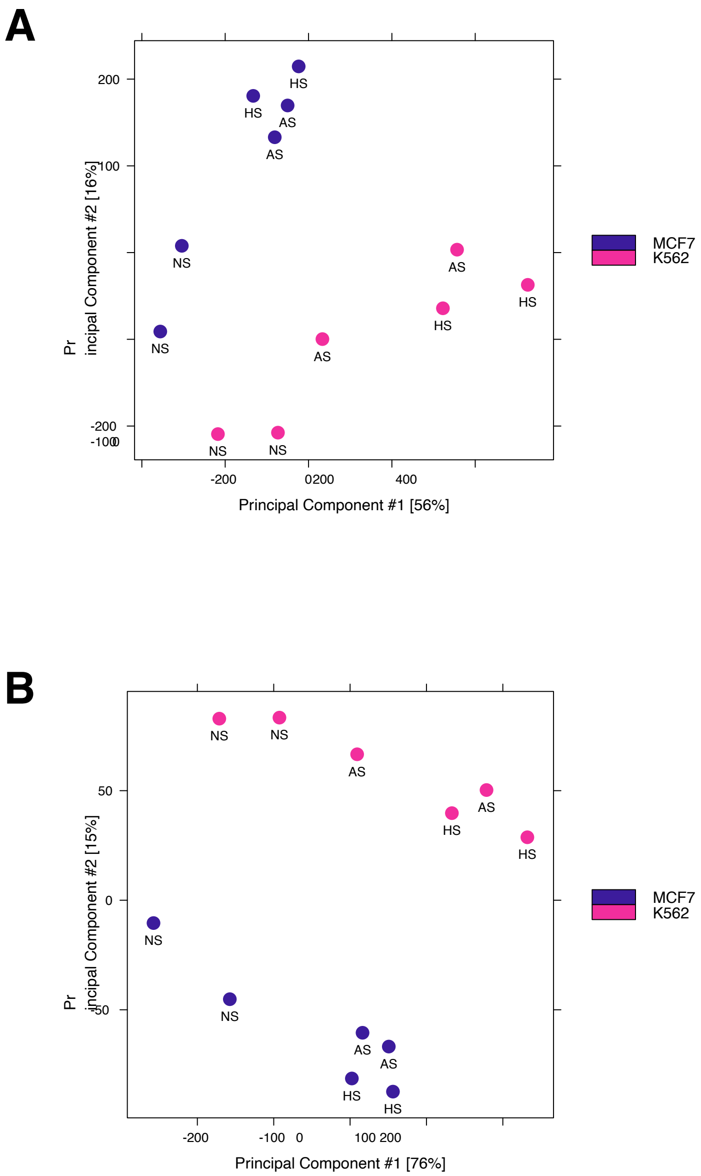


**Figure S10. Principal component analysis PCA plots for all HSF1 peaks**. The plots were made with diffbind using data after genomics copy-number correction wherein HSF1 signal for each peak region was normalized to the genomic copy number in the respective cell line (A). A diffbind PCA plot using a subset of HSF1 peaks that were common for both cell lines and treatments (B).

### Reference hg19

| Rank | Motif | Name | P-value | log P-pvalue | q-value (Benjamini) | # Target Sequences with Motif | % of Targets Sequences with Motif | # Background Sequences with Motif | % of Background Sequences with Motif |
|---|---|---|---|---|---|---|---|---|---|
| 1 | 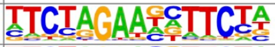 | AT3G09735(S1Falike)/col-AT3G09735-DAP-Seq(GSE60143)/Homer | 1e-1340 | -3.087e+03 | 0.0000 | 1598.0 | 58.15% | 2166.3 | 4.67% |
| 2 | 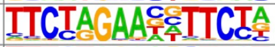 | HRE(HSF)/Striatum-HSF1-ChIP-Seq(GSE38000)/Homer | 1e-1319 | -3.038e+03 | 0.0000 | 1285.0 | 46.76% | 1024.9 | 2.21% |
| 3 | 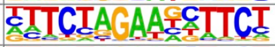 | HSF3(HSF)/colamp-HSF3-DAP-Seq(GSE60143)/Homer | 1e-1235 | -2.844e+03 | 0.0000 | 1957.0 | 71.22% | 4893.3 | 10.55% |
| 4 | 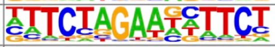 | HSFA6B(HSF)/colamp-HSFA6B-DAP-Seq(GSE60143)/Homer | 1e-1212 | -2.791e+03 | 0.0000 | 1602.0 | 58.30% | 2648.5 | 5.71% |
| 5 | 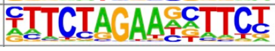 | HSFC1(HSF)/col-HSFC1-DAP-Seq(GSE60143)/Homer | 1e-1004 | -2.312e+03 | 0.0000 | 1089.0 | 39.63% | 1060.3 | 2.29% |

### K562 genome

| Rank | Motif | Name | P-value | log P-pvalue | q-value (Benjamini) | # Target Sequences with Motif | % of Targets Sequences with Motif | # Background Sequences with Motif | % of Background Sequences with Motif |
|---|---|---|---|---|---|---|---|---|---|
| 1 | 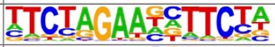 | AT3G09735(S1Falike)/col-AT3G09735-DAP-Seq(GSE60143)/Homer | 1e-1372 | -3.161e+03 | 0.0000 | 1598.0 | 58.15% | 2065.5 | 4.45% |
| 2 | 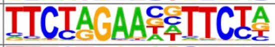 | HRE(HSF)/Striatum-HSF1-ChIP-Seq(GSE38000)/Homer | 1e-1326 | -3.055e+03 | 0.0000 | 1285.0 | 46.76% | 1010.5 | 2.18% |
| 3 | 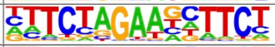 | HSF3(HSF)/colamp-HSF3-DAP-Seq(GSE60143)/Homer | 1e-1227 | -2.827e+03 | 0.0000 | 1957.0 | 71.22% | 4938.9 | 10.65% |
| 4 | 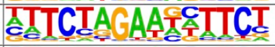 | HSFA6B(HSF)/colamp-HSFA6B-DAP-Seq(GSE60143)/Homer | 1e-1208 | -2.783e+03 | 0.0000 | 1602.0 | 58.30% | 2663.5 | 5.74% |
| 5 | 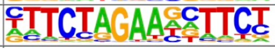 | HSFC1(HSF)/col-HSFC1-DAP-Seq(GSE60143)/Homer | 1e-1015 | -2.339e+03 | 0.0000 | 1089.0 | 39.63% | 1033.6 | 2.23% |

**Figure S11. Cell line specificity of HSF1 binding is unlikely to be affected by mutations.** The homer known motif output showing five topmost enriched motifs for peaks switching quartiles between MCF7 and K562 quartiles between cell lines (from Q1 in MCF7 to Q4 in K562 cells). Tables show representation of motifs for hg19 reference genome and K562-based single-nucleotide variants [33]. No quartile switching peaks were located within deletions specific to K562 as compared to hg19 reference.

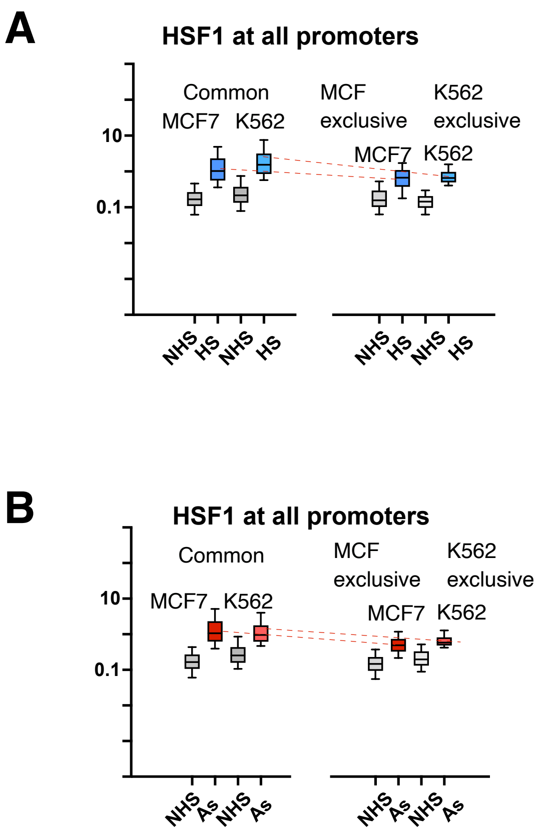


**Figure S12. HSF1 signal for peaks common and unique between cell lines.** Box plots for HSF1 signal are shown for MCF7 and K562 peaks that are common or unique for each cell line for HS (A) and As (B) treatments, for each biological replicate. Dashed lines indicate common and exclusive peaks for each cell line for the same replicate. The difference in HSF1 signal between these peaks is significant ($p<0.0001$) based on Mann-Whitney test.

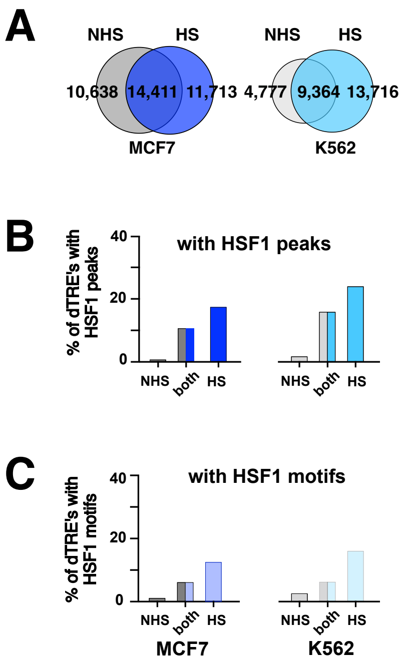


**Figure S13. Specificity of dTRE identification.** A. The overlap between dTREs called in NHS and HS conditions for each cell line. B. Percentage of dTREs with HSF1 peaks in NHS and HS treatments for each cell line. C. Percentage of dTREs with HRE sequence motifs for each cell line in NHS and HS treatments.

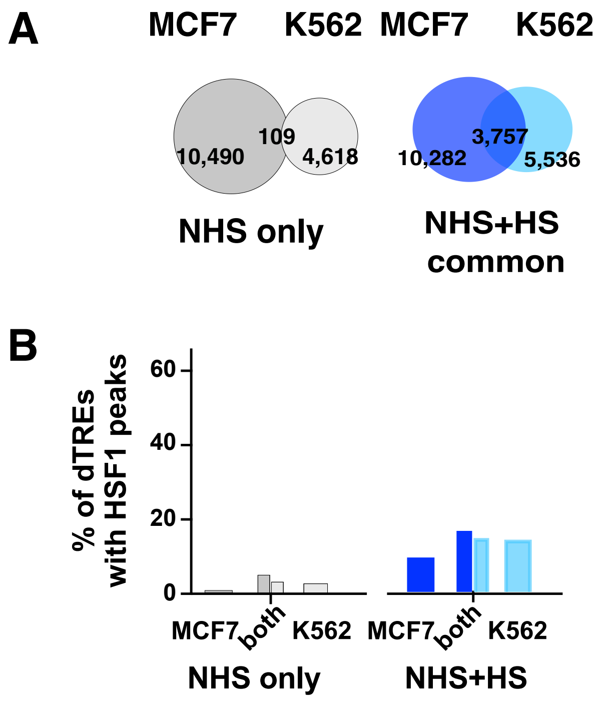


**Figure S14. Commonality of dTREs between distant cell lines.** A. The overlap between NHS only and NHS and HS common dTREs between MCF7 and K562 cell lines. B. The percentage of dTREs with HSF1 ChIP-seq peaks for the overlaps in (A).

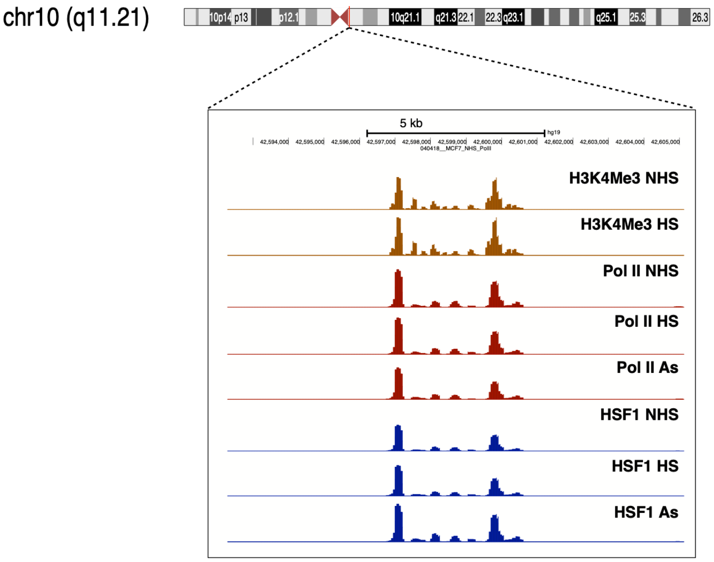


**Figure S15. An example of high-intensity peak found in basal (NHS) cells.** A UCSC genome browser track showing MCF7 cells with ChIP-seq datasets for a randomly selected high-intensity HSF1 peak. The location of the magnified region on the chromosome (centromere) is shown at the top.

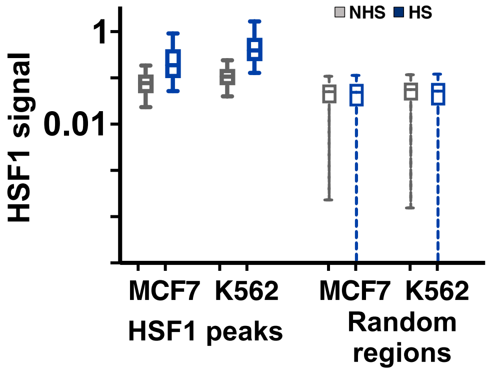


**Figure S16. NHS signal compared to the random background.** The box plot shows HS-only peaks in NHS and HS cells for each cell line as indicated (left) and signal from the same number of randomly chosen locations in the same datasets (right). The plots show means with error bars indicating 10-90% intervals.

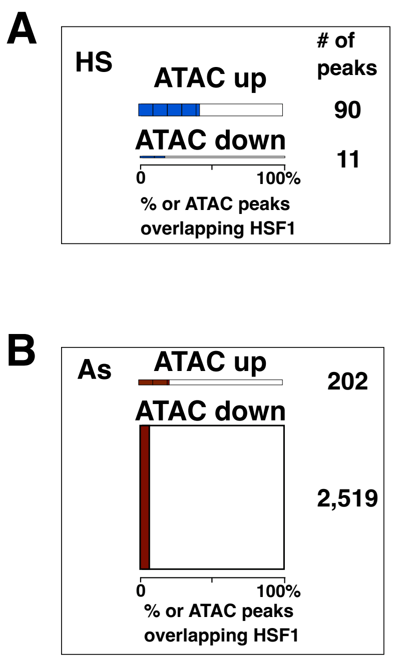


**Figure S17. Enrichment of HSF1 binding among up and downregulated ATAC regions.** The graphs show the number of up- and down-regulated ATAC peaks for HS (A) and As (B) treatments in MCF7 cells. The height of the bars is scaled to the number of peaks in each category (the numbers of peaks indicated on the right). The percentage of HSF1 peaks in each ATAC peak category is shown in X axis with blue (HS) and red (As) colors.

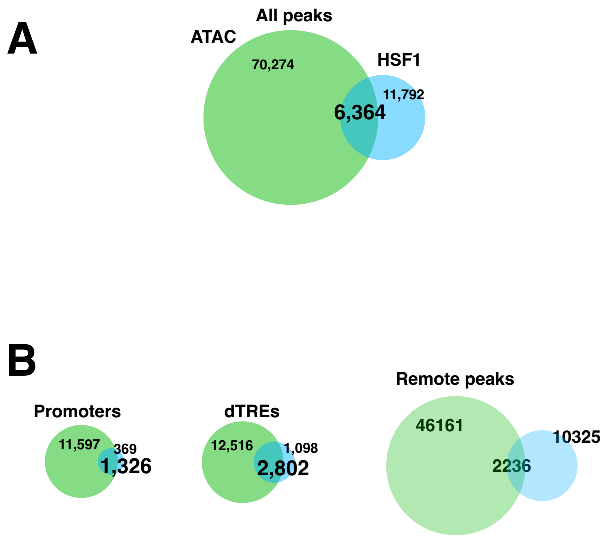


**Figure S18. Overlap between ATAC and HSF1 peaks.** A. Venn diagram showing the overlap between all ATAC and all HSF1 peaks in HS MCF7 cells. B. Venn diagrams showing the overlaps between peaks located in the indicated categories of genomic elements: promoters (left), dTREs (middle), and the rest of the genome (remote peaks, right).

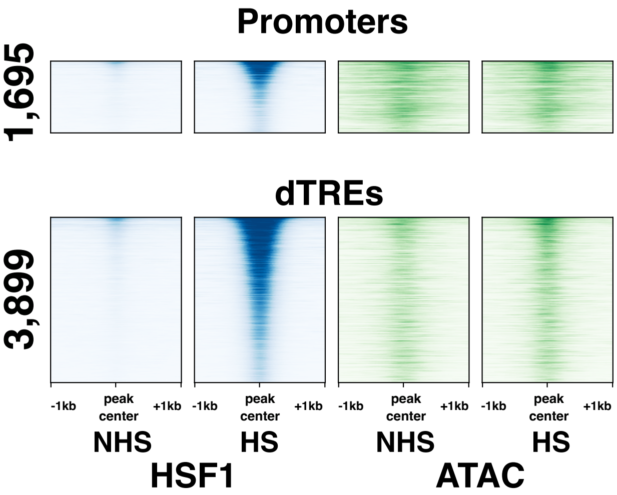


**Figure S19. HSF1 and ATAC signal in functional genomic elements.** HSF1 is binding to already open genomic elements. NHS and HS datapoints are shown for MCF7 cells for elements that overlap with HSF1 peaks including promoters (top) and dTREs (bottom). The numbers of peaks in each category is shown on the left of each heatmap.

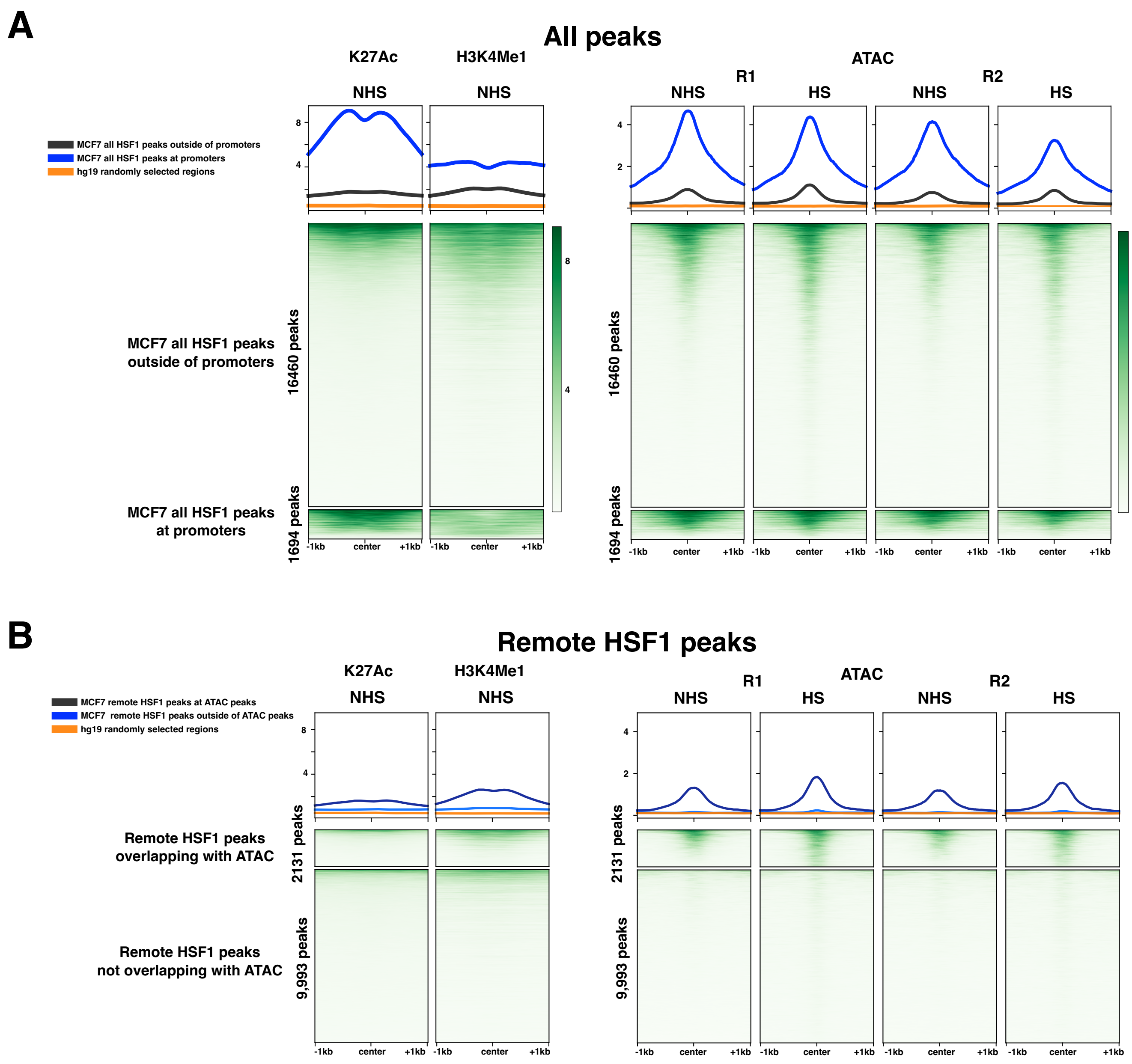


**Figure S20. ATAC and active histone marks around HSF1 peaks.** Indicated tracks are shown for all ATAC peaks overlapping with HS HSF1 peaks in MCF7 cells (A). B. The same tracks shown for remote HSF1 peaks for those that overlap with ATAC peaks (top) or have no overlapping ATAC peaks. Histone MCF7 cell tracks are derived from GSE85158 datasets.

### No overlap with ATAC

| Rank | Motif | Name | P-value | log P-pvalue | q-value (Benjamini) | # Target Sequences with Motif | % of Targets Sequences with Motif | # Background Sequences with Motif | % of Background Sequences with Motif |
|---|---|---|---|---|---|---|---|---|---|
| 1 | 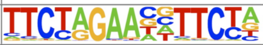 | HRE(HSF)/Striatum-HSF1-ChIP-Seq(GSE38000)/Homer | 1e-10564 | -2.433e+04 | 0.0000 | 8790.0 | 87.95% | 1659.2 | 4.16% |
| 2 | 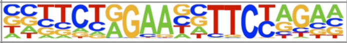 | HRE(HSF)/HepG2-HSF1-ChIP-Seq(GSE31477)/Homer | 1e-9504 | -2.188e+04 | 0.0000 | 7251.0 | 72.55% | 875.1 | 2.20% |
| 3 | 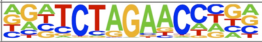 | ZBTB12(Zf)/HEK293-ZBTB12.GFP-ChIP-Seq(GSE58341)/Homer | 1e-925 | -2.132e+03 | 0.0000 | 2898.0 | 29.00% | 2883.0 | 7.23% |
| 4 | 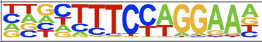 | Bcl6(Zf)/Liver-Bcl6-ChIP-Seq(GSE31578)/Homer | 1e-198 | -4.560e+02 | 0.0000 | 3982.0 | 39.84% | 10380.2 | 26.04% |
| 5 | 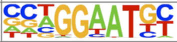 | TEAD4(TEA)/Tropoblast-Tead4-ChIP-Seq(GSE37350)/Homer | 1e-163 | -3.755e+02 | 0.0000 | 2765.0 | 27.67% | 6677.5 | 16.75% |

### Overlap with ATAC

| Rank | Motif | Name | P-value | log P-pvalue | q-value (Benjamini) | # Target Sequences with Motif | % of Targets Sequences with Motif | # Background Sequences with Motif | % of Background Sequences with Motif |
|---|---|---|---|---|---|---|---|---|---|
| 1 | 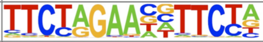 | HRE(HSF)/Striatum-HSF1-ChIP-Seq(GSE38000)/Homer | 1e-1200 | -2.764e+03 | 0.0000 | 1317.0 | 61.77% | 2058.5 | 4.31% |
| 2 | 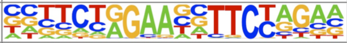 | HRE(HSF)/HepG2-HSF1-ChIP-Seq(GSE31477)/Homer | 1e-843 | -1.943e+03 | 0.0000 | 924.0 | 43.34% | 1252.2 | 2.62% |
| 3 | 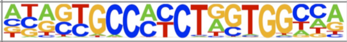 | CTCF(Zf)/CD4+-CTCF-ChIP-Seq(Barski_et_al.)/Homer | 1e-177 | -4.083e+02 | 0.0000 | 280.0 | 13.13% | 637.1 | 1.33% |
| 4 | 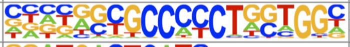 | BORIS(Zf)/K562-CTCFL-ChIP-Seq(GSE32465)/Homer | 1e-132 | -3.052e+02 | 0.0000 | 286.0 | 13.41% | 1012.3 | 2.12% |
| 5 | 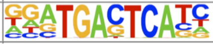 | Fra2(bZIP)/Striatum-Fra2-ChIP-Seq(GSE43429)/Homer | 1e-132 | -3.052e+02 | 0.0000 | 577.0 | 27.06% | 4199.2 | 8.78% |

**Figure S21. Sequence motifs among distant open and closed HSF1 peaks.** HSF1 peaks outside of promoters, dTREs, and annotated enhancers were separated into those that do and those that do not overlap with ATAC peaks, followed by homer motif identification. The first five most highly enriched motifs from homer's known motifs are shown for each group of HSF1 peaks. HSF1 peaks that are closed based on the absence of ATAC peaks show higher % of targets with HRE elements than those that are more open, that is, overlap with ATAC peaks.
